# Evolutionary and cellular analysis of the dark pseudokinase PSKH2

**DOI:** 10.1101/2022.09.10.507278

**Authors:** Dominic P Byrne, Safal Shrestha, Leonard A Daly, Vanessa Marensi, Krithika Ramakrishnan, Claire E Eyers, Natarajan Kannan, Patrick A Eyers

## Abstract

Pseudokinases, so named because they lack one or more conserved canonical amino acids that define their catalytically-active relatives, have evolved a variety of biological functions in both prokaryotic and eukaryotic organisms. Human PSKH2 is closely related to the canonical kinase PSKH1, which maps to the CAMK family of protein kinases. Primates encode PSKH2 in the form of a pseudokinase, which is predicted to be catalytically inactive due to loss of the invariant catalytic Asp residue. Although the biological role(s) of vertebrate PSKH2 proteins remains unclear, we previously identified species-level adaptions in PSKH2 that have led to the appearance of kinase or pseudokinase variants in vertebrate genomes alongside a canonical PSKH1 paralog. In this paper we confirm that, as predicted, PSKH2 lacks detectable protein phosphotransferase activity, and exploit structural informatics, biochemistry and cellular proteomics to begin to characterise vertebrate PSKH2 orthologues. AlphaFold 2- based structural analysis predicts functional roles for both the PSKH2 N- and C- regions that flank the pseudokinase domain core, and cellular truncation analysis confirms that the N-terminal domain, which contains a conserved myristoylation site, is required for both stable human PSKH2 expression and localisation to a membrane-rich subcellular fraction containing mitochondrial proteins. Using mass spectrometry-based proteomics, we confirm that human PSKH2 is part of a cellular mitochondrial protein network, and that its expression is regulated through client-status within the HSP90/Cdc37 molecular chaperone system. HSP90 interactions are mediated through binding to the PSKH2 C-terminal tail, leading us to predict that this region might act as both a *cis* and *trans* regulatory element, driving outputs linked to the PSKH2 pseudokinase domain that are important for functional signalling.

## Introduction

Protein kinases and pseudokinases are critical rate-limiting modulators of many aspects of signalling in both health and disease [1]. Pseudokinases are defined as lacking at least one of the conventional amino acids originally defined in active/canonical kinases, which coordinate ATP and/or facilitate transfer of a phosphate group from ATP to a protein substrate [2]. Historically, pseudokinase research has received only a fraction of the attention compared to the highly-detailed investigations of catalytically-active kinases, despite considerable evidence that conformational switching and subcellular scaffolding, as opposed to catalytic output *per se*, is fundamental for cellular signalling mediated by kinases and pseudokinases [3-6]. The unstudied (‘dark’) human pseudokinase PSKH2 [4, 7] is closely related to the Golgi-associated canonical kinase PSKH1 [8]. PSKH1 is a catalytically active member of the CAMK family that possesses a conventional autophosphorylating Ser/Thr kinase domain, a Golgi-targeting motif N-terminal to the kinase domain that is absent in PSKH2, and a putative Ca^2+^/CAM-dependent binding motif in the PSKH1 C-tail [2, 9]. A conserved regulatory C-tail is a common feature in kinases and pseudokinases, and sequence diversity in this region opens up the possibility for a variety of regulatory properties to be conferred [4, 10, 11]. PSKH2 proteins share significant sequence similarity with both PSKH1 and related CAMKs in the kinase domain, but diverge in the N and C-terminal segments flanking this region. Although a readily identifiable (‘unique’) PSKH2 gene is conserved across many vertebrate species, it displays species-specific variation including loss or retention of the catalytic Asp in the nominal kinase active site. The biology of PSKH2 remains uncharacterized at the molecular level, and our understanding of the enigmatic PSKH2 protein is currently restricted to inferred knowledge based on its amino acid sequence, predicted structure, and relation to PSKH1 (itself an understudied kinase). PSKH2 is a pseudokinase in primates due to a single Asp to Asn switch (N183) in the canonical HRD motif within the activation loop (Figure 1A). In protein kinases, the HRD Asp residue functions as a putative catalytic base critical for phosphoryl transfer to the -OH group of defined polar amino acids (Ser/Thr/Tyr) within a protein substrate. Lack of an Asp residue would therefore suggest that PSKH2 in primates is catalytically compromised. Other than the unusual HRN motif, all of the other amino acids residues typically required to coordinate ATP are conserved in PSKH2, and as such metal-dependent nucleotide-binding can be predicted to have been retained [4]. Interestingly, very low-level catalytic activity has been reported in analogous HRN containing pseudokinase domains of HER3 and JAK2 [12, 13], and it is possible that post-translational deamidation of Asn might also reinstate a functional Asp residue in this position, though no evidence exists for such a modification. However, PSKH2 in most non-primate higher chordates retains a conventional HRD motif [4], suggesting that the Asn-pseudokinase switch is a relatively recent evolutionary event. In addition to predicted abrogated kinase activity, PSKH2 deviates from PSKH1 in that the validated Golgi-targeting motif embedded in the N-terminal ‘unstructured’ domain of PSKH1 is conspicuously absent in PSKH2 [4, 8, 14]. This may suggest that both proteins occupy disparate spatial cellular niches. However, both proteins contain putative sites of N-terminal myristoylation and palmitoylation, a potential ‘smoking gun’ for membrane targeted proteins [4, 14].

**Figure 1.**
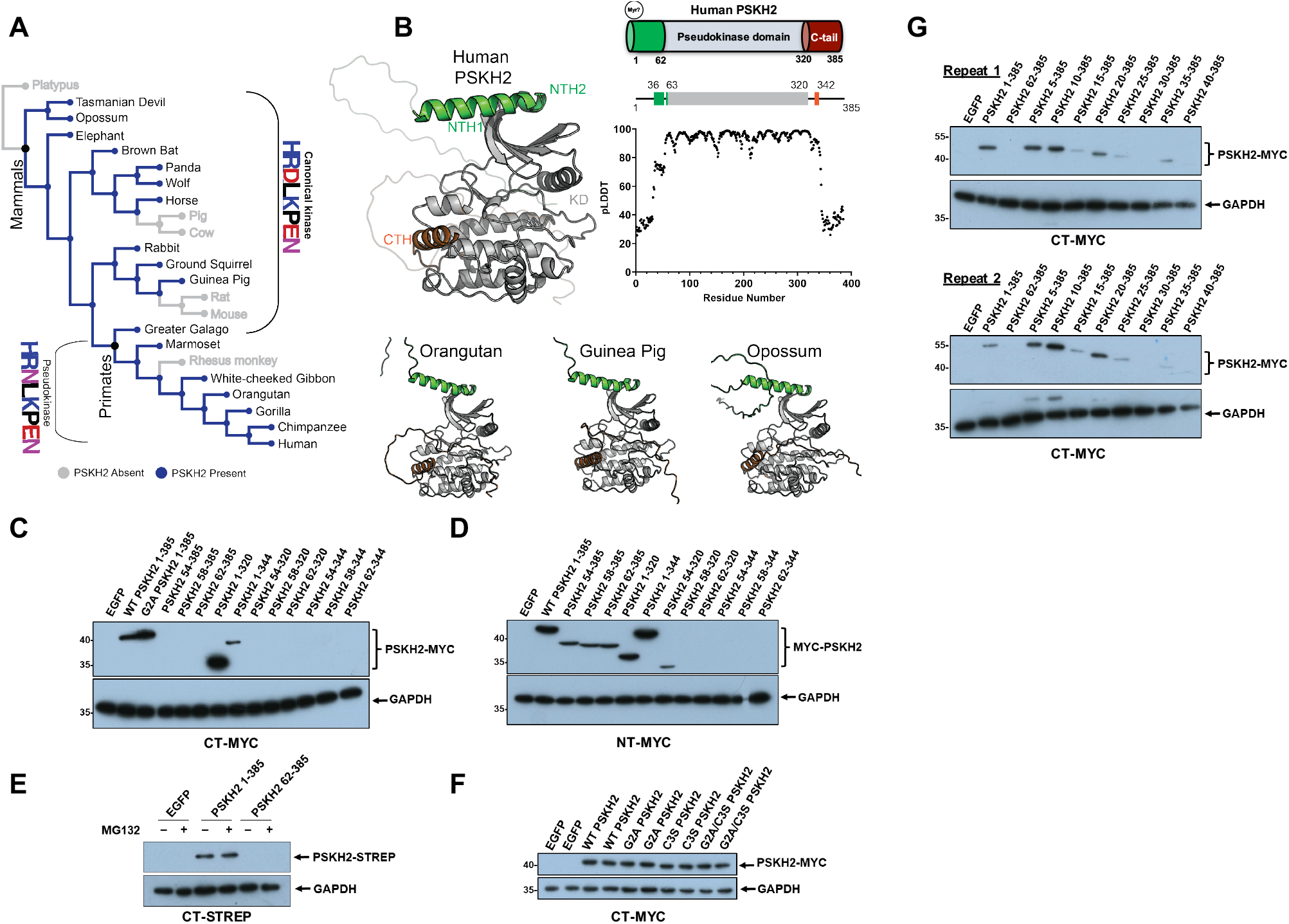
Informatics and Biochemical characterization of PSKH2. **(A)** Canonical and pseudokinase PSKH2 orthologs mapped to the SwissTree reference species tree. **(B)** Alphafold models for full length PSKH2 from Human (Q96QS6), Orangutan (H2PQQ4), Guinea Pig (H0VF24), and Opossum (F7ELA1). **(C)** Immunoblot showing expression of full length and truncated N-terminal (NT) or **(D)** C-terminal (CT)-MYC tagged PSKH2 variants in HEK-293T cells. **(E)** Expression of full-length (1-385) and truncated (62- 385) PSKH2 in the presence/absence of MG132. **(F)** Expression of WT PSKH2 and mutants containing amino acid substitutions at putative sites of N-terminal acylation. **(G)** Immunoblot showing changes in PSKH2 stability with incremental truncations at the N-terminus. Two independently repeated experiments are provided. All immunoblots constitute whole cell lysates processed with SDS-PAGE sample buffer. Transfection using a plasmid expressing EGFP was used as a control.

Phenotypic analysis of PSKH2 has been impeded by its absence from mouse and rat genomes, which prohibits standardized genetic knock-out approaches, and has also stymied biochemical characterization of the recombinant protein, which is highly challenging to express and purify in a soluble form [4]. Although we know little in regards to PSKH2 biological function, dys-regulation has also been associated with disease. For example, assessment of 24 cancer cohorts found significantly elevated copy numbers of *PSKH2* genes in 5-10 % of all patients [15] which was also prognostic for poor survival in some cancers [16]. Like many other ‘dark’ human kinases and pseudokinases, PSKH2 is included, but rarely discussed, in most high-throughput mRNA datasets (e.g. see https://maayanlab.cloud/Harmonizome/gene/PSKH2), which have been assembled from direct or indirect small molecule or genome-wide knock-down approaches in a variety of cell models [17, 18]. However, PSKH2 has been suggested to exhibit low-level synthetic lethality with the RAS oncogene [19], consistent with a role in proliferative signalling. PSKH2 is also a frequently mutated “dark” pseudokinase in human cancers, with the majority of mutations mapping to specific regions in the N-terminal domain and the pseudokinase domain [20].

In this study, we exploit evolutionary and organismal analysis [21, 22] and interface them with a new set of cellular tools to facilitate investigation of PSKH2. We find that vertebrate PSKH2 interacts with multiple cellular proteins, but itself lacks catalytic activity, regardless of the presence of a putative catalytic Asp residue. Cellular analysis of PSKH2 also establish that human PSKH2 requires an intact N-terminus for expression of the pseudokinase domain in cells. Alpha-fold models suggest an *in cis* interaction between the N-domain and the pseudokinase domain which we predict mediates folding and stabilisation, and mass-spectrometry-based proteomics demonstrates that human PSKH2 is enriched in a network of membrane-associated mitochondrial proteins. Moreover, the myristyolated N-terminus targets the protein to membrane-rich cellular fractions. Finally, we confirm that PSKH2 is a prominent client of the HSP90/Cdc37 chaperone system, and that this interaction is driven by the unique PSKH2 C-terminal tail, a putative Ca(2+)/CaM docking site in PSKH1 that may act as a common regulatory element, as previously established for a variety of AGC and CAMK kinases and pseudokinases.

## Results

### Evolutionary and structural analysis of PSKH2

To elucidate in-depth taxonomic information regarding canonical and pseudokinase PSKH2 orthologs, we leveraged the KinOrtho dataset [23] which contains 75 million kinase ortholog sequences across 17,000 proteomes (expanding ortholog counts by 30-60% in some families). For PSKH2 orthologs, we classified organisms containing either canonical (HRD-Asp) or pseudo (HRN-Asn) PSKH2 sequences and mapped the information onto the SwissTree species ‘tree’ [24]. While PSKH2 orthologs are present in a broad variety of mammals, it is only in primates that non-canonical pseudokinase PSKH2 sequences are found (Figure 1A).

AlphaFold 2 (AF2) is an artificial intelligence (AI) based protein structure prediction tool that can predict structures to near-experimental accuracy [25]. Due to the lack of any experimental structural data on PSKH2 (or the closely related PSKH1), we used AF2 to predict structures for both canonical and pseudokinase PSKH2 orthologs, and combined it with our knowledge of structural kinase studies. AF2 reports residue-specific confidence (pLDDT) on a scale from 0-100 [25]. For the human PSKH2 model (Figure 1B, Top Left), the kinase domain (residues 63-320) was modelled with medium-to-high confidence (pLDDT > 70) (Figure 1B, Top Right). However, most of the N (residues 1-35) and C (residues 344-385) termini regions were modelled with lower confidence (pLDDT < 70) (Figure 1B, Top). Interestingly, parts of the N-terminal regions are predicted to adopt α-helical conformations. Two putative N-terminal helices, termed helix 1 (NTH1) and helix 2 (NTH2), corresponding to residues 36-53 and 57-62 respectively in human PSKH2, as well as a single C-terminal helix (CTH), corresponding to residues 333-342, are predicted with medium-to-high confidence (Figure 1B). Moreover, helical regions in the flanking segments were consistently predicted, not only in humans, but also in all other vertebrate organisms (Figure 1B, Bottom).

While these non-pseudokinase domain regions are well conserved at the sequence level within PSKH2 orthologs, there are key differences in the flanking regions between PSKH1 and PSKH2 sequences (Supplementary Figures 1A, B). In PSKH2, two conserved prolines, Pro36 and Pro57 (Human PSKH2 numbering), initiate NTH1 and NTH2 (Supplementary Figure 1A). In contrast, PSKH1 sequences contain additional prolines between residues 69-86 (Human PSKH1 numbering, UniProt ID: P11801) (Supplementary Figure 1A) which results in a shorter NTH1 in the human PSKH1 model (Supplementary Figure 1C, D). In the C-terminal flanking region, conserved proline, Pro346, shortens the first helix (CTH) in PSKH2 compared with our PSKH1 model (Supplementary Figure 1B, C, D). Moreover, due to the presence of an additional helix in PSKH1, the mechanism whereby the C-terminal flanking region tethers to the kinase domain is distinct between PSKH1 and PSKH2 (Supplementary Figure 1C, D). The structural and functional roles of these flanking segments are explored in the following sections.

### Biochemical analysis of PSKH2 expression in human cells

Based on data deposited in the human protein atlas database (https://www.proteinatlas.org/), PSKH2 expression is classified as ‘extremely low’, or ‘absent’, in most well-studied laboratory cell lines, including HEK-293. In addition to a lack of specific biochemical tools, such as validated antibodies or small molecules, this has impeded characterisation of PSKH2 protein function in the endogenous cellular context. To improve our biochemical understanding of PSKH2, we initially generated a transient overexpression system in HEK-293T cells, generating PSKH2 constructs with 3C protease cleavable N- or C-terminal MYC tags (Figure 1C & 1D). In addition to full length PSKH2, we also truncated the protein at the N- or C terminus of the pseudokinase domain. Interestingly, regardless of the position of the affinity tag, removal of the N-terminal region was very poorly tolerated and resulted in a loss of PSKH2 expression, based on immunoblotting (Figure 1C & 1D). In contrast, C-terminally truncated PSKH2 proteins were relatively stable. Expression of N-terminally truncated PSKH2 could not be rescued by the inclusion of the proteasome inhibitor MG132, suggesting that the observed loss of pseudokinase expression was not a consequence of targeted proteasomal degradation (Figure 1E). Furthermore, we did not detect an observable loss of PSKH2 following mutation of the predicted putative sites of myristoylation (Gly 2) and/or palmitoylation (Cys 3), indicating that these post-translational modifications (PTMs) are not the critical determinants for protein stability observed in the N-terminal truncated forms of the protein (Figure 1F). This conclusion is supported by the observation that inclusion of an N-terminal fusion tag, which blocks co-translational myristoylation of proteins [26], did not reduce expression of full-length PSKH2 proteins.

Inspection of the AF2-predicted structure of PSKH2 (Figure 1B) suggests that the kinase domain adopts a typical bi-lobal conformation, and is flanked by N- and C-terminal tail regions that are largely disordered. The transition between disordered and order in the flanking regions of kinases and pseudokinases is of broad potential interest, since it provides a potential mechanism for regulation of protein:protein interactions. As previously discussed, our model predicts an extra alpha helix structural element (NTH1, between residues 36-53) that was omitted in all of the N-terminally truncated constructs tested (Figure 1B, Supplementary Figure 1A). Consistently, incremental truncation of the N-terminus of PSKH2 resulted in a loss of PSKH2 expression following deletion of the first ∼25 amino acids, whereas smaller truncations N-terminal of the potential helix had minimal impact (Figure 1G).

To maintain a functional PSKH2 N-terminus with an intact, modifiable, myristoylation site at Gly2, we affinity purified full length PSKH2 with a cleavable, tandem C-terminal Strep tag, and subjected it to tandem analysis MS (Supplementary Figure 2A), in order to study PSKH2 post-translational modifications (PTMs). Following proteolytic elution of PSKH2 by 3C-mediated cleavage of the affinity tag, the presence of PSKH2 was confirmed using an in-house generated polyclonal PSKH2 antibody that recognises overexpressed PSKH2 (Supplementary Figure 2B). We further verified the specificity of this PSKH2 antibody in HEK-293T expressing PSKH2-STREP, and *E. coli* cells induced to express (denatured) full-length 6His-PSKH2 (Supplementary Figure 2C-D). In HEK-293T cells, PSKH2 and STREP-tag reactive bands were detected at the same molecular weights, confirming the presence of tagged-PSKH2, and these signals were absent in control cells transfected with EGFP (Supplementary Figure 2B). Interestingly, the PSKH2 antibody also detected a doublet with a higher electrophoretic mobility than the predicted exogenous PSKH2 band in both transfection conditions, which could suggest the co-purification of endogenous PSKH2, or perhaps more likely, promiscuous immunoreactivity with another unidentified protein.

LC-MS/MS analysis of either tryptic, chymotryptic or elastase-generated peptides from affinity purified PSKH2 expressed in HEK-293 cells yielded a sequence coverage of ∼63 % considering all three proteolytic enzymes. Given the identification of PSKH2 as a member of the human kinome, we also subjected these peptides to phosphopeptide-enrichment prior to LC-MS/MS analysis to search for phosphorylation sites across the PSKH2 polypeptide (Supplementary Figure 2A). Although 6 phosphorylation sites have been catalogued in the PhosphoSite Plus repository (https://www.phosphosite.org/proteinAction.action?id=2123&showAllSites=true), to our knowledge, none have been identified in more than a single high-throughput (shotgun) proteomics study, and we were unable to detect these sites of modification in any of our MS runs. However, we consistently observed phosphorylation at Tyr 228 in the pseudokinase domain using this approach. We subjected immunoprecipitated human PSKH2 to immunoblotting analysis to probe for total Ser/Thr and Try phosphorylation. While we observed a faint signal indicative of Tyr phosphorylation at approximately the size of PSKH2, we did not observe a signal with a generic pSer/Thr antibody (Supplementary Figure 2E), consistent with the notion that under these experimental conditions, full-length PSKH2 is not highly-modified by phosphorylation.

A primary goal of our study was to isolate sufficient quantities of PSKH2 to determine whether it retains the capability to bind to ATP. Unfortunately, obtaining sufficient yields of pure PSKH2 was challenging, prohibiting effective analysis of this aspect of PSKH2 functionality. However, an *in vitro* kinase assay using immunoprecipitated PSKH2 in the presence of either ^32^P-ATP or ATP-γ-S suggested that human PSKH2 was incapable of autophosphorylation (Figure 2, Supplementary Figure 3A). Introduction of a predicted ‘inactivating’ mutation of the canonical metal-binding residue in the DFG motif, D204A, had no detectable effect on the very low levels of phosphotransferase activity present, suggesting the small amount of detectable Tyr phosphorylation was most likely due to phosphorylation by an endogenous kinase in cells. Interestingly, restoration of the catalytic HRD motif (N183D) in human PSKH2 was also insufficient to activate the pseudokinase in terms of autophosphorylation under these experimental conditions. Moreover, kinase assays using immunoprecipitated PSKH2 proteins from a variety of vertebrate species also demonstrated that naturally occurring HRD and HRN variants are all inactive under these conditions (Supplementary Figure 3B-C). This could either suggest that all PSKH2 is inactive after isolation regardless of the catalytic apparatus in each species, that additional co-factors (such as Ca^2+^/CaM, which target the PSKH1 C-terminal region [8]) are required for activity and are absent in our immunoprecipitations, or that it is autoinhibited through an unknown mechanism. An additional possibility is that, despite a lack of autophosphorylation, immunoprecipitated PSKH2 possesses activity towards a physiological substrate missing from the assembled reaction. In order to test this hypothesis broadly, we incubated PSKH2 isolated from human cells with common non-specific substrates, including myelin basic protein (MBP), α-casein and a selection of histone proteins. However, we were unable to detect robust PSKH2-dependent phosphorylation of any generic substrates (Figure 2B), though our controls demonstrate that the assays would have detected these events. Interestingly PSKH1 was also shown to lack phosphorylation of non-specific substrates *in vitro* [8], suggesting that substrate phosphorylation is either linked to a very specific substrate (as demonstrated for the Mg^2+^-independent pseudokinase CASK and its substrate neurexin [27, 28], or is inactive in the context of the full-length polypeptide under these conditions.

**Figure 2.**
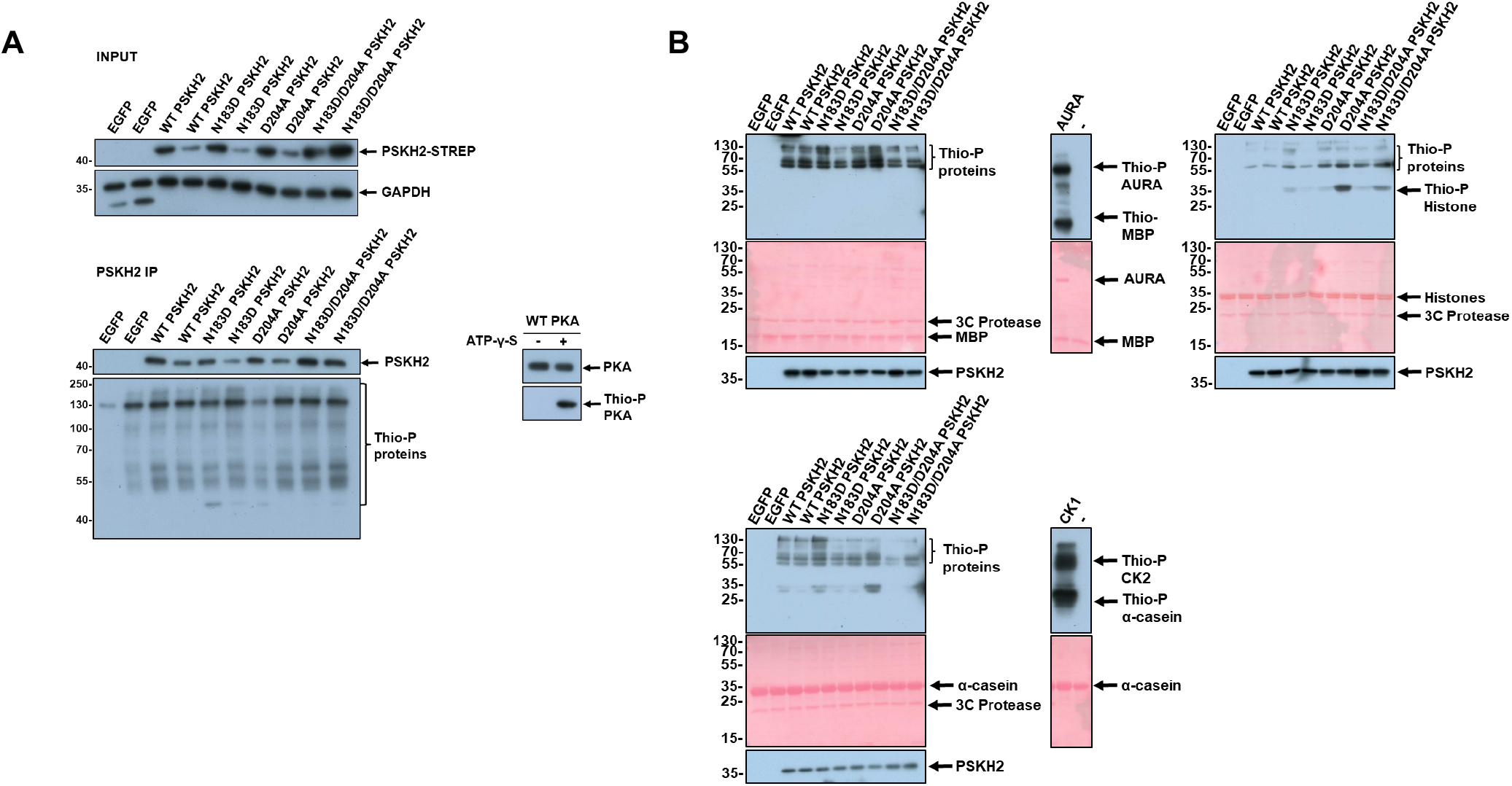
Analysis of kinase activity associated with PSKH2 immunoprecipitates. **(A)** Kinase assays were performed containing material precipitating with PSKH2 from HEK-293T cells in the presence of ATP-γ-S. Protein phosphorylation as a consequence of kinase activity was detected using an antibody with specificity towards thiophosphate esters (Thio-P), and is compared to precipitations from cells overexpressing EGFP, WT PSKH2, or ‘canonical’ amino acid changing variants of PSKH2 in the absence or **(B)** presence of the indicated polypeptide substrate. PKA, Aurora A (AURA) and casein kinase 2 (CK2) were used as positive controls for auto- and substrate phosphorylation.

### Proteomic analysis of the human PSKH2 interactome

To gain new insights into the cellular functions of PSKH2, we evaluated the PSKH2 interactome in HEK-293 cells using LC-MS/MS. N- and C-terminal STREP-tagged PSKH2 were isolated from HEK-293T cells in triplicate, and physically-eluted from Strep-TACTIN beads using a 3C protease treatment. PSKH2 contains predicted putative sites for myristoylation and palmitoylation at the second Gly and third Cys residues respectively, which are conserved in PSKH1, and are molecular determinants for localisation to the Golgi membrane [14]. Given that PSKH2 might therefore be membrane associated, our affinity precipitation protocol was performed in the presence of the detergent n-Dodecyl-β-D-Maltoside (DDM). Over 170 proteins were identified in the complement of binding partners observed in at least two biological replicates for N- and C-terminally tagged soluble PSKH2. These were all statistically-enriched compared to control immunoprecipitation conditions [p-value > 0.01]) (Figure 3A, Supplementary table 1). Strikingly, we observed discrepancies in the interaction networks of N- and C-terminal tagged PSKH2 (Figure 3B, Supplementary Figure 4A and B, Supplementary table 2 and 3), suggesting that these flanking regions might serve specific protein-protein interaction functions. A second interpretation of this finding is that the presence of an affinity tag might change the PSKH2 interactome. Particularly noteworthy was UNC119, a myristoyl-binding protein required to coordinate the trafficking of other myristoylated proteins [29], which was only observed in immunoprecipitates containing C-terminally tagged PSKH2. This presence of an N-terminal affinity tag blocks co-translational myristoylation of expressed proteins, which is therefore predicted to preclude precipitation and thus identification of myristoyl-binding proteins. Several interactions that have previously been reported in broad interactome network analysis of the human kinome, including HSP90, CDC37, RABGGTB, UNC119 and UNC119B [30] were also found to associate with PSKH2 in our analysis. We also showed that RABGGTB could be detected by immunoblotting analysis of affinity-purified PSKH2-containing samples (Supplementary Figure 5C), confirming that it as a component of an PSKH2 interaction network.

**Figure 3.**
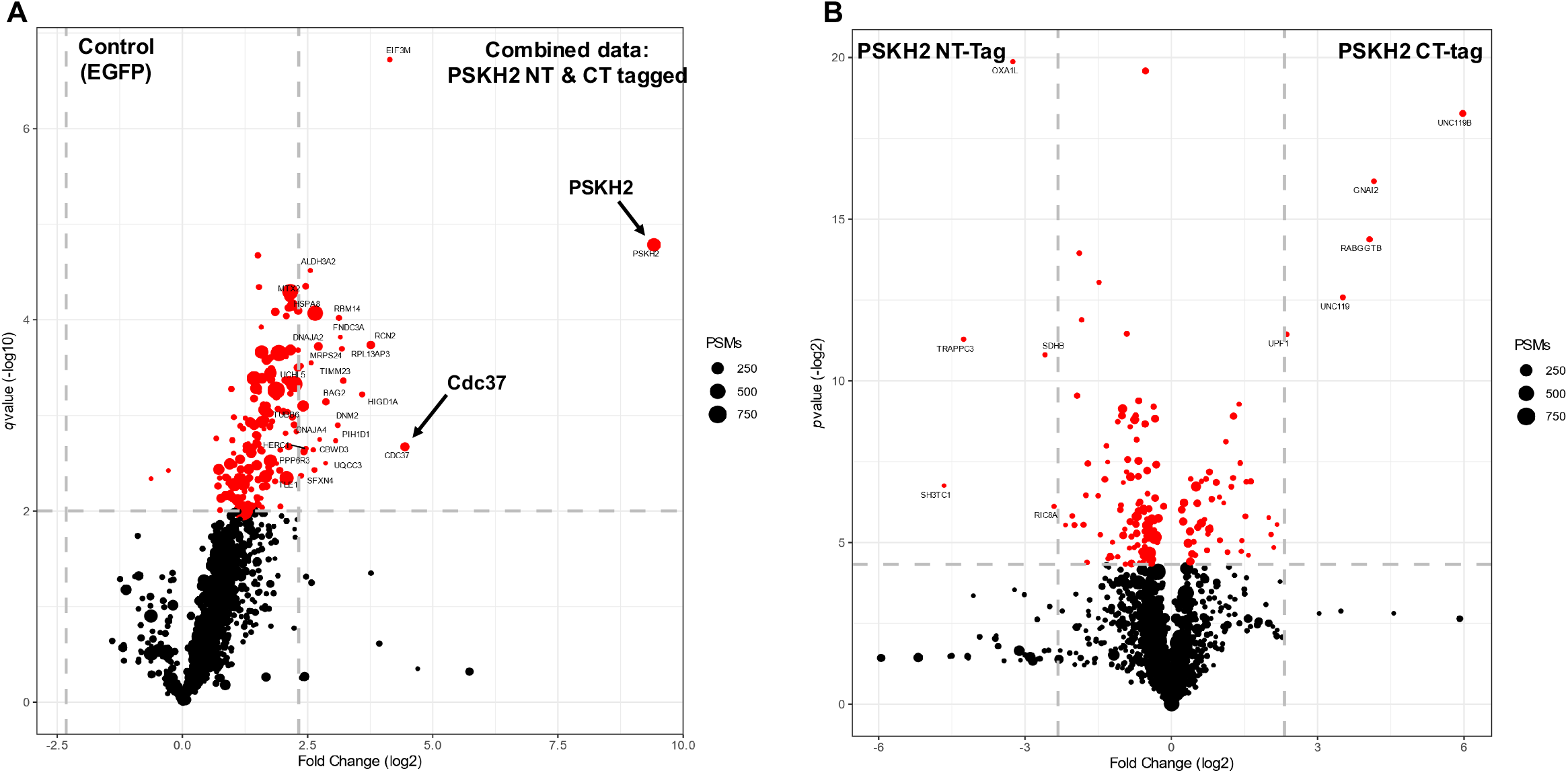
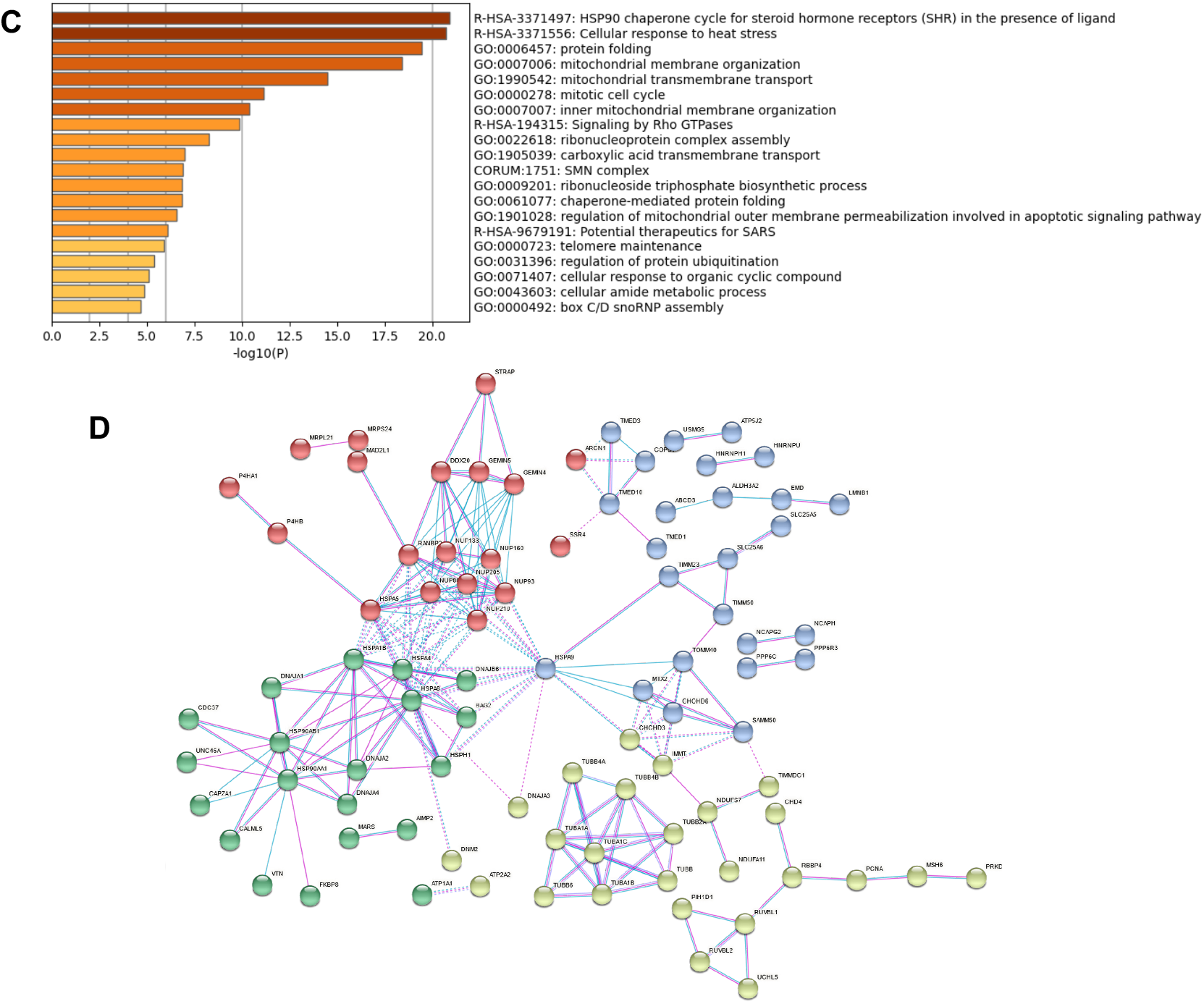
Analysis of the PSKH2 protein interactome. **(A)** Volcano plot depicting label-free protein quantification from PSKH2 IPs, shown as −log10(*q* value) versus log2 (1/21% abundance fold change). Dot size equates to the combined number of MS/MS events for a protein across all replicates. Black, *q* value > 0.01; red, *q* value < 0.01; significant proteins with a comparative fold change >5 are labelled with their gene name. Gray-dashed lines denote fivefold change and *q* value = 0.01. Data shown represents changes in abundance of proteins co-immunoprecipitated with both NT- and CT-tagged PSKH2 compared to an EGFP-transfected control experiment and **(B)** comparative abundance changes of proteins between NT- and CT-tagged PSKH2 IPs conditions. For both comparisons, only proteins observed in at least two replicate IPs were analysed. **(C)** GO term enrichment analysis of interactors that were significantly elevated compared to EGFP controls and present in both NT and CT-tagged PSKH2 IPs (*p* value < 0.01) using Metascape. **(D)** Protein-protein interaction networks for significant binding partners (*p* value < 0.01) in both NT and CT-tagged PSKH2 IPs.

We next subjected our PSKH2 interactome data sets to functional enrichment analysis using Metascape [31]. The most significantly enriched functional ontology terms for PSKH2 binding partners included HSP90 chaperones, the cellular response to heat stress, protein folding, mitochondrial membrane organization, the mitotic cell cycle, and inner mitochondrial membrane organization (Figure 3C). Protein-protein interaction enrichment analysis of the data set also identified several specific clusters of known interacting proteins (Figure 3D, Supplementary Figure 5A and B). These densely-connected networks of proteins included clusters of mitochondrial-associated import and resident proteins (TIMM23, TIMM50, TOMM40, HSPA9, SLC25A6), chaperones and co-chaperones involved in kinase and non-kinase protein folding (HSP90, CDC25C, HSPA1B, DNAJA1,2 & 4, HSPA8) and nuclear associated proteins (NUP160, NUP210, RANBP3). Due to a lack of validated PSKH2 antibodies, confident expression of endogenous PSKH2 is absent in most human tissues. Where it has been analyzed as part of the human cell atlas assembly, PSKH2 has been mapped to the nucleoplasm [32, 33], and when tagged with an N-terminal FLAG sequence, PSKH2 (and PSKH1) have previously been described as ‘cytosolic’ or ‘nuclear’. However, the use of NT-FLAG epitope tags for both PSKH1 and PSKH2 in these studies is likely to interfere with N-terminal myristoylation, which has the potential to abolish physiological cellular interaction networks. As we show below, N-terminal modification(s) are important for PSKH2 subcellular distribution, conceptually similar to findings with PSKH1 and its targeting to the Golgi apparatus [14]. Informatics analysis also identified molecular links between PSKH2 and specific networks of nuclear proteins, which supports the previous assertion (based on immunofluorescence-based analysis of tagged PSKH2) that PSKH2 might function as a nuclear-associated protein. Most surprisingly however, was a very strong connection between PSKH2 and proteins of validated mitochondrial origin, and to our knowledge this is first time that such a relationship has been seen for human PSKH2. The prominence of binding-partners that are frequently recruited to membranous organelles and predicted putative PSKH2 myristoylation strongly suggests that PSKH2 is a membrane-associated protein, similar to other network members such as those connected to Rho GTPase signalling.

### PSKH2 is a client of the HSP90 chaperone system

One of the most significantly enriched proteins in immunoprecipitation studies with PSKH2 was Cdc37 (Supplementary Table 1). Cdc37 was also previously identified as a PSKH2-binding partner in untargeted proteome-wide interactome studies [30, 34] and is one of the few interactors captured by commercial BioGRID software. Cdc37 is a molecular chaperone component of the HSP90 complex, which is of particular significance given its role in facilitating the correct folding of multiple members of the human kinome, including CDK4, CDK6, SRC, RAF-1, MOK and AKT [35-37]. HSP90α and HSP90β were also enriched in our affinity-capture MS analysis (Supplementary Table 1). Our finding confirms previous investigations that have established an interaction between PSKH2 and endogenous HSP90 [38]. We further confirmed the interaction between PSKH2 and HSP90 (and two other HSP90 partner proteins, Cdc37 and FKBP5) by immunoblotting following affinity purification of either N- or C-terminal Strep-tagged PSKH2 (Figure 4A). In addition, PSKH2-Strep could be co-immunoprecipitated using FLAG-tagged variants of HSP90, Cdc37 or FKBP5 as bait (Supplementary Figure 6). Next, we immunoprecipitated PSKH2 or truncated variants of PSKH2 with a C-terminal MYC epitope. All three HSP90 complex proteins were detected following immunoprecipitation of PSKH2 with MYC affinity resin, and truncation of the N-terminus or substitution of the putative site of myristoylation had no discernible effect on HSP90-binding (Figure 4B, Supplementary. Figure 7A). In marked contrast, deletion of the PSKH2 C-terminal tail (Figure 1), completely eliminated HSP90-complex protein interactions, suggesting that the interaction between PSKH2, HSP90 and its cognate co-chaperones is directed through this region. Finally, we interrogated the effect of ATP on PSKH2 complex formation. For this purpose, we performed scanning mutational analysis at canonical ATP binding positions within the kinase domain of PSKH2 in order to generate constructs predicted to be deficient in ATP binding, specifically K92H and D204A (analogous to the canonical Lys72 and Asp184 residues in PKA involved in ATP coordination). In addition, we re-introduced a catalytic Asp at position Asn183 (N183D) to more closely resemble a typical canonical protein kinase. Compared to wild-type human PSKH2 protein, none of the mutations had an appreciable effect on HSP90 binding (Figure 4C, Supplementary Figure 7B). This implies that ATP binding is dispensable for engagement with the HSP90 complex in human PSKH2. Interestingly, HSP90 complex proteins could also be co-precipitated with PSKH1 and several other mammalian PSKH2 variants (Supplementary Figure 7C & D), confirming conservation of an interaction. Together, these data present further evidence that PSKH2 (and PSKH1) are both clients of the HSP90 chaperone complex, an interaction that may be required to promote appropriate folding or complex formation, or to prevent misfolding in cells. Experimental analysis of the biochemical properties of PSKH2 has long been stymied by the instability of the protein and the poor quality of preparations obtained using traditional protein purification techniques. A functional requirement for chaperone-assisted folding may partly explain this phenomenon. In order to investigate this hypothesis further, HEK-293T cells overexpressing C-terminal tagged PSKH2 were exposed to increasing concentrations of the HSP90 inhibitor geldanamycin. Geldanamycin exerts anti-HSP90 activity by competitively inhibiting HSP90 ATPase activity, resulting in the misfolding and degradation of client proteins [39-41]. At the highest geldanamycin concentrations tested, PSKH2 levels were diminished by ∼70 % compared to control (DMSO) treatments (Figure 4D, Supplementary Figure 8A), whereas the concentrations of GAPDH, tubulin, Cdc37 and HSP90 were unperturbed. Importantly, levels of AKT (a known HSP90 target) were also reduced (but not eliminated) under the same conditions (Figure 4D, Supplementary Figure 8A). Consistently, treatment of cells with celastrol, a compound that specifically disrupts formation of the HSP90/Cdc37 complex [42], also resulted in a marked depletion of PSKH2 in cells (Figure 4E, Supplementary Figure 8B). Finally, exposure to dexamethosone to stimulate upregulated expression of the HSP90 co-chaperone and PSKH2 binding partner, FKBP5 [43, 44] resulted in a consistent ∼2-fold increase in PSKH2 levels (Supplementary Figure 8C). Unexpectedly, the synthetic geldanamycin derivative and HSP90 inhibitor, tanepsimycin (17-AAG) [45], increased the expression of exogenous PSKH2 in HEK-293T cells (Supplementary Figure 8D). The reason for this discrepancy is unclear, but could be a consequence of an off-target effect of the compound, possibly related to cell stress and the induction of apoptosis [46].

**Figure 4.**
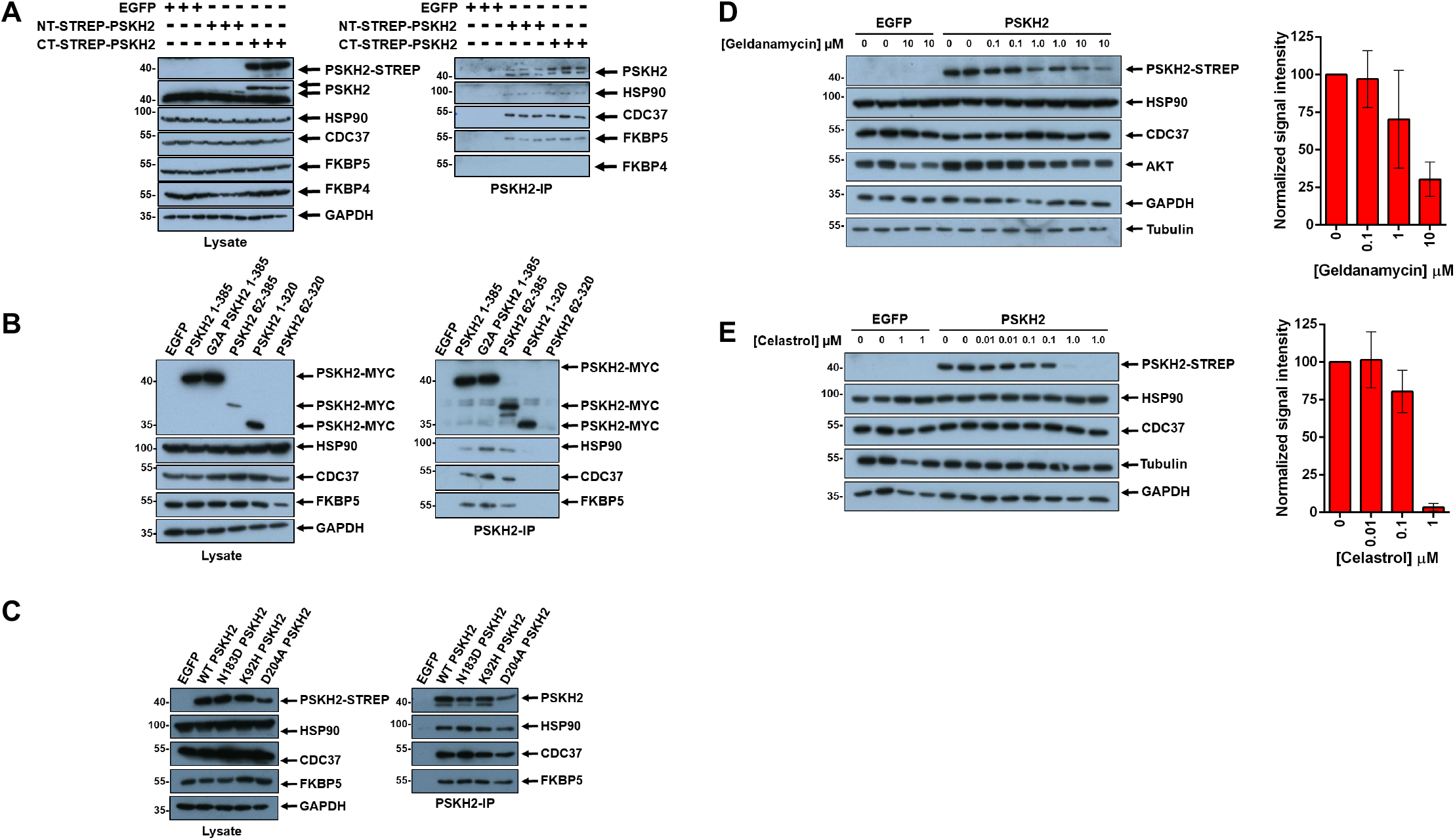
PSKH2 is a client of the HSP90 chaperone system. **(A)** Co-precipitation of HSP90, CDC37 and FKBP5 with NT- and CT-strep tagged PSKH2. **(B)** Co-precipitation of HSP90 proteins with full-length and truncated PSKH2 constructs. **(C)** Co-precipitation of HSP90 proteins with WT and canonical kinase activity mutants of PSKH2. **(D)** Reduction of PSKH2 expression in HEK-293T cells with increasing concentration of geldanamycin and **(E)** celasterol. Densitometry data shown in the right panel of (D) and (E) is mean and SD of PSKH2 band intensity from four independent experiments, first normalized to tubulin and then PSKH2 bands corresponding to control (DMSO) conditions.

### Molecular dynamics analysis of human PSKH2

To build on our finding that PSKH2 is a bona fide client of the HSP90 system through its C-tail region, we analysed the intrinsic dynamics of different human PSKH2 sub-domains *in silico*. Initially, we performed 700 ns long molecular dynamics (MD) simulation using the AF2 model of full length PSKH2. This revealed major differences in the interaction of the N and C terminal flanking regions with the pseudokinase domain. Mobile parts of the protein can be identified by calculating the Root Mean Square Fluctuations (RMSF) of the residues during the course of the simulation. The N-terminal flanking region bordering the pseudokinase domain, specifically residues 27-37, were highly mobile, and residues 350-365 in the C-terminal tail (Figure 1A) were also mobile when compared to the rest of the protein (Supplementary Figure 9A). Interestingly, the N-terminal region dislodges from the kinase domain within the first 10 ns, whereas the C-terminal flanking region maintains its interaction with the pseudokinase domain throughout the simulation (Supplementary Figure 9B, Supplementary Video 1). The interaction of the C terminal flanking region with the pseudokinase domain is stabilized by both salt bridges and hydrophobic interactions throughout the MD simulation (Supplementary Figure 9B). Notably, Trp369, a residue within the C-terminal flanking region that is only present in PSKH2, (Supplementary Figure 1) is buried within the hydrophobic pocket created by Phe146 from the αD helix, Pro186 from the catalytic loop, Ala254 from the αF helix, and Leu260 from a loop between the αF and αG helices (Supplementary Figure 9B). Moreover, Arg375 forms salt bridge with Glu100 and Glu103 from αC helix interchangeably (Supplementary Figure 9B). Additional hydrophobic interactions are observed between terminal Leu381, Leu384, and Leu385 residues in the C terminal region, with Leu234 and Leu235 in the region following the activation loop, as well as Leu275 and Tyr271 from the αG helix (Supplementary Figure 9B). Despite being modelled with relatively low confidence by AlphaFold (Figure 1B), the C terminal flanking region might therefore stabilize the pseudokinase domain when not engaged with the HSP90 complex, performing a regulatory in cis function, analogous to those reported for other CAMK and AGC kinases [10].

Using AlphaFold-multimer, an AF-based prediction tool that can be used to model higher order homo- and hetero-complexes [47], we next predicted putative structural interactions between Cdc37, HSP90A/B and full-length PSKH2. Compared to the PSKH2 model alone (Figure 1B), the predicted structure of PSKH2 in a complex with Cdc37 and HSP90A/B reveals additional helices in both terminal regions (Supplementary Figure 9C). Notably, only the C-terminal helices are predicted to interact with both Cdc37 and HSP90A/B (Supplementary Figure 9C, Supplementary Video 2), supporting our experimental finding that revealed the PSKH2 C-tail as the key determinant for interaction with the HSP90 complex in cells.

### PSKH2 participates in a network enriched in mitochondrial proteins

As reported above, clusters of proteins associated with the mitochondria, including TIMM23 (a translocase of the inner mitochondrial membrane), co-purify with human PSKH2 expressed in human cells. Multiple mechanisms are recognised for the import of mitochondrial proteins [48]. We confirmed a physical interaction between PSKH2 and endogenous TIMM23 after immunoprecipitation (Figure 5A, Supplementary Figure 10A). Next, we probed for PSKH2 in various subcellular fractions prepared by centrifugation of HEK-293T cell extracts overexpressing C-terminal MYC tagged PSKH2. Consistently, PSKH2 was enriched in preparations that contained the mitochondrial marker protein ATP5A (Figure 5B), and from which predominantly cytoplasmic proteins such as GAPDH and EGFP were excluded. Interestingly, we were unable to detect enrichment of PSKH2 in nuclear-enriched extracts (Supplementary Figure 10B).

**Figure 5.**
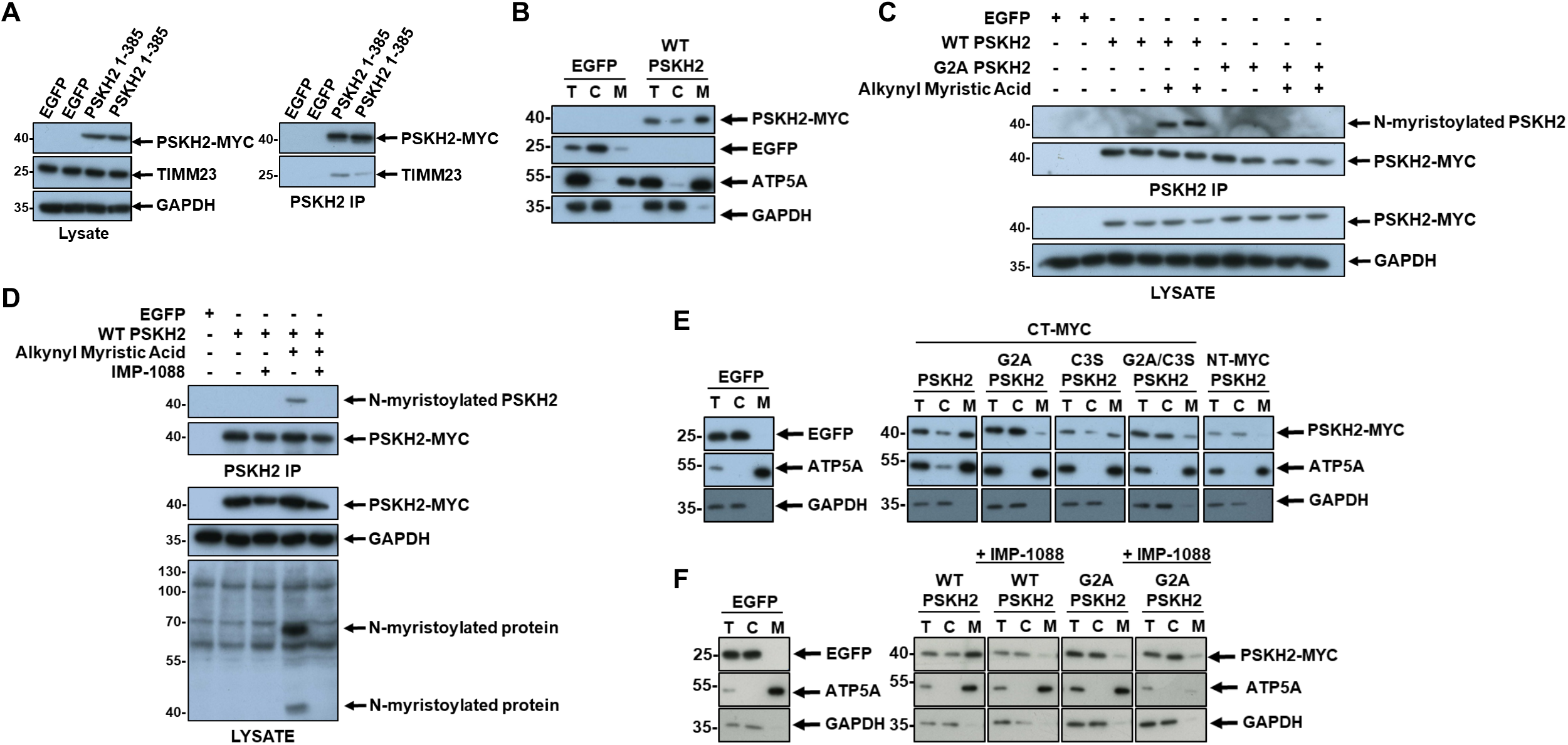
Detection of PSKH2 in membrane-enriched fractions is dependent on N-myristoylation. **(A)** Co-precipitation of mitochondrial protein TIMM23 with PSKH2. **(B)** Mitochondrial fractionation of HEK-293T cells overexpressing EGFP of CT-Myc PSKH2. **(C)** Immunoprecipitation of PSKH2 from HEK-293T cells metabolically labelled with alkynyl myristic acid. Myristoylated protein was labelled with biotin azide using a copper-catalyzed click reaction and detected after western blotting using neutravidin-HRP. **(D)** Detection of myristoylated PSKH2 in immunoprecipitations from HEK-293T cells treated in the absence of presence of IMP-1088. **(E)** Mitochondrial fractionation of HEK-293T cells overexpressing EGFP, WT PSKH2 (CT-Myc) or PSKH2 mutated at putative sites of acylation. **(F)** Mitochondrial fractionation of PSKH2-expressing HEK-293T treated in the presence or absence of IMP-1088. For all fractionations; T = total cell lysate, C = cytoplasmic fraction, M = mitochondrial-enriched fraction.

We next sought to elucidate the mechanism(s) through which PSKH2 is recruited to this mitochondria-rich fraction. For some proteins, such as the kinase PINK1 (PTEN-induced putative kinase 1), localisation to the mitochondrial membrane can be directed through a short N-terminal mitochondrial targeting sequence [49], although we were unable to identify such a sequence in human PSKH2. However, in the absence of a canonical targeting sequence, post-translational myristoylation has been shown to support mitochondrial association of proteins such as TOMM40 and SAMM50 [50], and the canonical protein kinase AMPK [51]. Despite careful interrogation of our MS data, we were unable to detect peptides from the N-terminal region of PSKH2 under any conditions, which prohibits assessment of acylation at either of the putative myristoylation and palmitoylation sites [4]. We next set out to study the potential for acylation in PSKH2, using a variety of approaches. Initially, HEK-293T cells transiently overexpressing C-terminal MYC-tagged PSKH2 or an acylation blocking G2A control, were incubated for 18 h in the presence or absence of the ‘clickable’ reagent alkynyl myristic acid. PSKH2 was then immunoprecipitated from whole cell lysates, and click chemistry employed to conjugate biotin-azide to labelled PSKH2, which was then evaluated using a neutravidin-HRP reporter. Following cellular myristate labelling, only isolated WT PSKH2 incorporated the alkynyl derivative, whilst myristoylation was absent in a G2A mutant (Figure 5C). These results demonstrate that PSKH2 is likely to be myristoylated in cells at the Gly2 position on the N-terminal domain. In support of this finding, exposure of cells to IMP-1088, a potent dual inhibitor of human *N*-myristoyltransferases NMT1 and NMT2 [52] eliminated N-myristoylation of PSKH2 (Figure 5D). These observations are also consistent with our earlier co-affinity purification analysis, which revealed an interaction with CT-tagged PSKH2 and UNC119B, a myristoyl-binding protein [29].

To evaluate the role of myristoylation in the recruitment of PSKH2 to mitochondrial fractions, we collected subcellular extracts from HEK-293T cells overexpressing WT, G2A, C3S or G2A/C3S CT-MYC tagged PSKH2. Mutation of the myristoylated Gly alone was sufficient to abrogate PSKH2 signal in mitochondrial-enriched fractions, resulting in a marked increase of the protein in the cytosol (Figure 5E, Supplementary Figure 10C). In contrast, substitution of the predicted palmitoylated cysteine (C3S) did not induce pronounced redistribution of PSKH2. Similarly, modification of PSKH2 with a N-terminal MYC tag (which blocks co-translational myristoylation) generated PSKH2 that was almost exclusively localised within the cytoplasmic fraction (Figure 5E). Finally, we exposed cells to IMP-1088 and evaluated biochemical localisation in extracts. Inhibition of total cellular *N*-myristoyltransferase activity efficiently blocked targeting of PSKH2 to the mitochondrial fraction as effectively as elimination of the Gly residue (Figure 5F, Supplementary Figure 10 D-E). As a control, we did not observe a reciprocal loss of mitochondrial ATP5A (which is not known to be myristoylated) under identical experimental conditions. Based on three repeat experiments, PSKH2 signal intensity was decreased in the mitochondrial-rich membrane fraction by ∼ 65 % in the presence of IMP-1088. Interestingly, under all tested experimental conditions, PSKH2 was also detected in cytoplasmic fractions, suggesting a dynamic shuttling mechanism for PSKH2 in cells.

## Discussion

Of the most understudied human kinases and pseudokinases, PSKH2 remains the most enigmatic, and no small molecules or validated cellular phenotypes are currently available with which to interrogate its function. This claim is supported by the fact that, to our knowledge, there is not a single publication in the literature where PSKH2 is the primary investigative focus, and as such, virtually every facet of PSKH2 biochemistry remains obscure. This lack of data is likely a consequence of multiple factors, not least the fact that this pseudokinase is challenging to prepare and analyse in recombinant form, which has prevented structural, biochemical or standard small molecule screening approaches [53-55]. Moreover, PSKH2 protein is detected at extremely low abundance in most tissues. Finally, PSKH2 has not been functionally associated with human disease as a ‘driver’, and its absence from mice and rat genomes has prevented conventional genomic and phenotypic characterisation. In order to begin to address these issues, we have undertaken a broad informatics and cell-based analysis of this unusual human pseudokinase.

Our attempts to generate soluble full-length or truncated human PSKH2 in bacteria, including the use of a variety of solubility enhancing affinity tags, bacterial expression strains and induction conditions, have all been unsuccessful. As such, despite PSKH2 retaining many of the canonical catalytic residues that are needed for canonical kinases (and some pseudokinases) to bind to ATP in a metal-dependent manner, a biochemical appraisal of its properties has not been possible due to a lack of folded material. PSKH2 instability in the context of bacterial expression likely reflects a requirement for protein co-factors and PTMs to facilitate correct folding and/or stabilisation. Indeed, although we were able to express and partially purify (immunoprecipitate) PSKH2 from HEK-293T cells, expression was highly sensitive to conserved sequences in the N and C-tails regions, and this stability might also be dependent on interaction(s) with HSP90 and the co-chaperones Cdc37 and FKBP5 (Figure 4). The chaperone-assisted folding and maturation of many kinases and pseudokinases has previously been established, although the biological role of these interactions remains obscure in many cases [35, 36]. However, there remains a clear role for molecular chaperones during the generation of many biologically-active kinases, including those prone to unfolding [56]. The specificity determinants that characterise HSP90 binding interfaces within client proteins are often defined by stretches of hydrophobic amino acids with a net-positive charge [57, 58]. Distal to the kinase domain, the C-terminal region of PSKH2 (320-385) is predicted to be intrinsically disordered (Figure 1B), with an amino acid composition consisting of ∼37 % hydrophobic and ∼22 % basic residues. Interestingly, compared to the theoretical isoelectric points (pI) of the N-terminal region and kinase domain (pI of 8.9 and 8.6 respectively), the PSKH2 C-terminal tail exhibits a compositional bias towards basic amino acids, with an estimated pI of ∼12.5 and a net charge of *z* = + 8.71 at physiological pH (*z* of the N-terminus and kinase domain are + 1.2 and + 3.6 respectively). These characteristics may dispose the PSKH2 C-tail to accommodate HSP90 binding and stabilise the protein in cells [56]. Indeed, PSKH2 lacking an intact C-terminal tail is completely deficient in HSP90-binding (Figure 4). However, the removal of the C-terminal tail has markedly less effect on PSKH2 stability compared to truncations within the N-terminus, which suggests that interplay between HSP90 and PSKH2 instead functions to modulate the largely unstructured C-tail, which we therefore predict could function as both an *in cis* and *in trans* docking element in cells. In addition to stabilisation of cellular PSKH2, it is also tempting to speculate that an interaction with HSP90 may also be regulatory; for example Cdc37-dependent HSP90 binding locks CDK4 in a semi-folded and inactive conformation [37, 59]. However, the lack of a clear physiological role for PSKH2 makes assignment of a similar role for HSP90 challenging, although we speculate that it may be relevant in the context of other regulatory proteins that PSKH2 colludes with in cells, the identify of which remain to be established. As an aside, recombinant human PSKH2 co-expression in bacteria with a combination of molecular chaperones (DnaJ, DnaK, GroEL, GroEZ, GrpE, and Tig) was also insufficient to resolve stability issues associated with the human pseudokinase.

### Role of the PSKH2 N-terminal region

We identified the N-terminal PSKH2 region as a crucial stability determinant for the pseudokinase domain, since truncation of this region was poorly tolerated in cells. Although a functional role for the N-terminal region is yet to be assigned, it is interesting that this largely disordered region is potentially structurally significant for PSKH2. In addition to containing a predicted alpha helix (of unknown function), the N-terminus of PSKH2 is predicted by informatics to be co- and post-translationally acylated [4]. Interestingly, specific mutational analysis demonstrates that acylation status has no bearing on PSKH2 stability in cells (Figure 1F). We do, however, find experimental evidence for an extreme N-Gly-myristoylation site, which facilitates subcellular localisation of PSKH2 to mitochondrially-enriched subcellular fractions (Figure 5). Physiological myristoylation is known for several human protein kinases, including the Ser/Thr kinase PKA [60] and the tyrosine kinase ABL [61]. Mitochondrial recruitment of PSKH2 was abrogated by mutating the site of myristoylation, or by directly inhibiting NMT1/2 enzyme activity in cells. Insertion of the hydrophobic myristoyl chain into a lipid bilayer is important for the interaction of a swathe of myristoylated enzymes to membranes, where signaling can be driven through substrate proximity [62]. Enzymatic attachment of a myristoyl group can also regulate protein functionality, a phenomenon known as a ‘myristoyl switch’ [61, 62]. In this regard, it is noteworthy that subcellular localisation of the closely related PSKH1 kinase to the Golgi-membrane is also driven by a myristoylation-dependent mechanism [4, 8, 14], as is mitochondrial recruitment of the related canonical protein kinase AMPK [51]. For PSKH1, and other membrane-associated proteins, a dual ‘myristoyl-palmitoyl’ switch is known to be required to promote membrane localisation [8, 14, 63, 64]. Although amino acids for myristoylation and palmitoylation are conserved in both PSKH2 and PSKH1 across species, only the former modification appears to be required for enrichment of human PSKH2 in mitochondrial containing fractions (Figure 5E), since substitution of the putative site of S-palmitoylation (Cys 3) had no effect on the amount of co-purifying PSKH2. Interestingly, affinity-capture MS-based sequence analysis of the PSKH2 N-terminal domain was consistently hampered in our hands by incomplete coverage, even when employing multiple proteases. Direct detection and site determination of native protein/peptide palmitoylation by MS is notoriously challenging due to the instability of the modification during sample preparation and its loss during tandem MS [65]. More work is required to validate modification of Cys3 or distal cysteines by S-palmitoylation. However, Geranylgeranyl transferase type-2 subunit beta (RABGGTB) was also enriched in IPs of CT-tagged PSKH2, and this suggest that PSKH2 could also be a target for isoprenylation [66]. Further experimentation is needed to validate and clarify the regulatory roles of these modifications, and to determine the extent to which lipid modifications regulate PSKH2 localisation and function. Since prokaryotes do not possess N-myristoyl transferase enzymes, this also further explains confounding issues with the expression of full-length N-myristoylated proteins such as PSKH1 and PSKH2 in bacterial systems [67].

We have yet to uncover a clear biological role for the PSKH2 pseudokinase domain in any species. However, in this study we generated experimental evidence that PSKH2 is co-translationally N-myristoylated and that this modification is required for recruitment of the protein to a mitochondrial-rich subcellular fraction. Myristoylation can control enzyme activation *in cis*, as well as concentrating signaling components on membranes [68]. Using affinity capture MS, we detected interactions between PSKH2 and multiple mitochondrial proteins, which fully supports our biochemical fractionation analysis. Regulatory roles for several kinases in mitochondrial homeostasis have previously been reported. For example, association of AMPK with damaged mitochondria is sufficient to promote selective mitophagy and cell survival [69], PINK1 plays a prominent role in mitochondrial life and death [70], and Aurora A possesses cryptic mitochondrial import encoded by proteolytically-processed amino acids in the non-kinase N-terminal 35 amino acids [71]. Further studies are needed to tease apart the roles of PSKH2 in human biology, including its potential interaction with the mitochondrial netork. These might include human cellular CRISPR ‘knock-in’ studies in different cell types, where the functional consequences of PSKH2 membrane association and HSP90 biology can be dissected in terms of cell signaling alongside cytological analysis of PSKH2 (and PSKH1) subcellular localisation; these important analyses are beyond the scope of the current work.

In this paper, we demonstrate that full-length vertebrate PSKH2 fully justifies its pseudokinase moniker [72, 73], since PSKH2 protein sequences from a variety of species lack detectable catalytic activity in terms of ATP-dependent substrate phosphorylation, regardless of the presence or absence of the putative catalytic Asp residue. Analysis of AF2 models representing a variety of vertebrate PSKH2 proteins leads us to propose an *in cis* interaction between an alpha helix in the N-domain and the pseudokinase domain. Mass spectrometry-based analysis of PSKH2 complexes demonstrates that the human pseudokinase is enriched in a network of membrane-associated mitochondrial proteins. Moreover, a myristoylated N-terminus targets PSKH2 to preparations of membrane-rich cellular fractions. We also demonstrate unequivocally that PSKH2 is a client of the HSP90/Cdc37 chaperone system, and that this interaction is driven by the unique PSKH2 C-terminal tail; based on structural modelling, we hypothesise that this HSP90-targeting region could act as a mobile regulatory element, as previously established for a variety of AGC and CAMK kinases and pseudokinases. Future work will build upon these findings, in order to reveal the biological role(s) of PSKH2, and the closely related PSKH1, which appeared after their recent evolutionary divergence in the vertebrates.

## Materials and Methods

### Reagents

General biochemicals, unless otherwise stated, were purchased from Sigma-Aldrich. Primers for molecular cloning and site directed mutagenesis were produced by integrated DNA technologies (IDT). Reagents for metabolic labelling of acylated PSKH2 were purchased from Click Chemistry Tools. ATP-γ-S, Thiophosphate ester antibody and *para*-nitrobenzyl mesylate were purchased from Abcam. All commercial antibodies were purchased from Cell Signalling Technology. A PSKH2 polyclonal antibody was raised toward a unique peptide in the C-lobe of the human PSKH2 pseudokinase domain (KGKYNYTGEPWPSISC) and affinity-purified prior to testing (Abgent).

### Human cell culture, immunoprecipitation, and Western blot analysis

HEK-293T cells were cultured in Dulbecco’s modified Eagle medium (Lonza) Supplementarylemented with 10% fetal bovine serum (HyClone), penicillin (50 U/ml), and streptomycin (0.25 μg/ml) (Lonza) and maintained at 37°C in 5% CO2 humidified atmosphere. All exogenously expressed proteins were cloned into a pcDNA3 vector and expressed in frame with a 3C protease cleavable, N- or C-terminal Myc, FLAG or tandem STREP tag. HEK-293T cells were transfected using a 3:1 polyethylenimine (PEI [branched average *M*w ∼25,000 Da; Sigma-Aldrich]) to DNA ratio (30:10 μg, for a single 10-cm culture dish). Point mutations were generated using standard PCR-based mutagenic procedures. For co-expression experiments, 5 μg DNA was used for each protein with 30 μg PEI. Whole cell lysates were collected 48 h post transfection in bromophenol blue–free SDS-PAGE sample buffer Supplementarylemented with 1% Triton X-100, protease inhibitor cocktail tablet, and phosphatase inhibitors (Roche), and sonicated briefly. Total cell lysates were clarified by centrifugation at 20,817*g* for 20 min at 4°C, and supernatants were sampled and diluted 30-fold for calculation of the protein concentration using the Coomassie Plus Staining Reagent (Bradford) Assay Kit (Thermo Fisher Scientific). Cell lysates were normalised for total protein concentration and processed for immunoblotting. Where indicated, cells were incubated with 10 µM of the proteasome inhibitor MG132, or an equivalent volume of DMSO vehicle control (0.1% v/v DMSO) for 4 h prior to harvesting the cells. Geldanamycin, celastrol, dexamethasone and 17-AAG were prepared in DMSO, and added to PSKH2 or mock-expressing HEK-293T cells 4 h after transfection. Cells were cultured in the presence of the indicated concentration of inhibitor (or 0.1% DMSO) for 18 h prior to collection of total cell lysates.

For immunoprecipitation experiments, proteins were harvested 48 h post transfection in a lysis buffer containing 50 mM Tris-HCl (pH 7.4), 150 mM NaCl, 0.1% (v/v) Triton X-100, 1 mM DTT, 1% (w/v) dodecyl-β-D-Maltoside (DDM), 0.1 mM ethylenediaminetetraacetic acid (EDTA), 0.1 mM ethylene glycol-bis(β-aminoethyl ether)-*N*,*N*,*N*′,*N*′-tetraacetic acid (EGTA) and 5% (v/v) glycerol and Supplementarylemented with a protease inhibitor cocktail tablet and a phosphatase inhibitor tablet (Roche). Lysates were briefly sonicated on ice and clarified by centrifuged at 20,817*g* for 20 min at 4°C, and the resulting supernatants were incubated with either Pierce Anti-c-Myc-Agarose resin (Thermo Fisher Scientific), Strep-TactinXT 4Flow (iba) or anti-FLAG G1 Affinity Resin (GeneScript) for 1-3 hours (as required) with gentle agitation at 4°C. Affinity beads containing bound protein were collected and washed three times in 50 mM Tris-HCl (pH 7.4) and 150 mM NaCl and then equilibrated in storage buffer (50 mM Tris-HCl [pH 7.4], 100 mM NaCl, 1 mM DTT, 1% (w/v) DDM and 5% (v/v) glycerol). More stringent washing (500 mM NaCl) was applied for kinase assays. The purified proteins were then eluted from the suspended beads over a 1-hour period with 3C protease (0.5 μg) at 4°C, with gentle agitation. Elution of FLAG-tagged HSP90, Cdc37 or FKBP5, and PSKH2 species in kinase assay was achieved using SDS-PAGE sample buffer.

### Affinity-capture Mass spectrometry

Affinity captured PSKH2-STREP samples were diluted 10-fold in 25 mM ammonium bicarbonate (pH 8.0) and reduced with dithiothreitol and alkylated with iodoacetamide [74], 0.2 μg trypsin gold (Promega) was added and incubating at 37°C with gentle agitation for 18 h. Digests were then subjected to strong cation exchange using in-house packed stage tips, as previously described [75]. Dried peptides were solubilized in 20 μl of 3% (v/v) acetonitrile and 0.1% (v/v) TFA in water, sonicated for 10 minutes, and centrifuged at 13,000 x *g* for 15 min at 4 °C prior to reversed-phase HPLC separation using an Ultimate3000 nano system (Dionex) over a 60-minute gradient [74]. All data acquisition was performed using a Thermo Orbitrap Fusion Lumos Tribrid mass spectrometer (Thermo Scientific), with higher-energy C-trap dissociation (HCD) fragmentation set at 32% normalized collision energy for 2+ to 5+ charge states. MS1 spectra were acquired in the Orbitrap (60K resolution at 200 *m/z*) over a range of 350 to 1400 *m/z*; AGC target = standard, maximum injection time = auto, with an intensity threshold for fragmentation of 2e^4^. MS2 spectra were acquired in the Iontrap set to rapid mode (15K resolution at 200 *m/z*), maximum injection time = 50 ms with a 1 min dynamic exclusion window applied with a 0.5 Da tolerance. For binding partner analysis, data was searched twice (search settings were identical, with the addition of low abundance resampling imputation of missing values in the 2^nd^ search) using Proteome Discoverer 2.4; searching the UniProt Human Reviewed database (updated weekly) with fixed modification = carbamidomethylation (C), variable modifications = oxidation (M), instrument type = electrospray ionization–Fourier-transform ion cyclotron resonance (ESI-FTICR), MS1 mass tolerance = 10 ppm, MS2 mass tolerance = 0.5 Da. Percolator and precursor ion quantifier nodes (Hi3 Label free Quantification (LFQ)) were both enabled. All data was filtered to a 1% False discovery rate. Data from the 1^st^ search was put into a custom R script that extracted all protein accessions with LFQ data in at least 2 of 3 replicates of a condition which was used to filter the imputation containing dataset. Accessions were used to obtain the gene name using the UniProt ID retrieval tool. Fold changes and T-tests were calculated and log2 transformed, before importing into a custom R script for plotting. For PSKH2 tag orientation plotting, LFQ data was normalized to the level of PSKH2 for that replicate prior to calculations. Data was additionally analyzed through PEAKS Studio (version XPro) using the same database, mass tolerances and modifications [75]. PEAKS specific search settings: instrument = Orbi-Trap, Fragmentation = HCD, acquisition = DDA, De Novo details = standard and a maximum of 5 variable PTMs possible. PEAKS PTM mode was enabled and filtering parameters of De Novo score >15, −log10P(value) >30.0, Ascore >30.0 and seen in at least 2 of 3 replicates were applied for a PTM to be maintained.

### *In vitro* kinase assays

15 µl affinity resin containing precipitated PSKH2 protein from a single transfected 10 cm culture dish was equilibrated in 50 mM Tris-HCl [pH 7.4], 100 mM NaCl, 1 mM DTT, 1% (w/v) DDM, 5% (v/v) glycerol, 10 mM MgCl2 and assayed in the presence of the indicated substrates and 1mM [γ^32^P] ATP (20 μCi ^32^P per assay) or ATP-γ-S for 1 h at 30°C with gentle agitation. [γ^32^P] ATP reactions were terminated by denaturation in SDS sample buffer. ATP-γ-S assays were incubated with 5 mM *para*-nitrobenzyl mesylate (*p*NBM) for a further 30 mins at 20°C to alkylate thiophosphorylated proteins prior to the addition of SDS sample buffer. Proteins were resolved by SDS-PAGE and analysed by autoradiography or western blotting with the appropriate antibody. Where appropriate, assays were also performed in the presence of 5 µg of the indicated polypeptide substrates.

### Subcellular fractionation and generation of mitochondrial-enriched preparations

HEK-293T cells were transfected with PSKH2-MYC or EGFP, and collected after 48 h. Cell pellets were resuspended in NKM buffer (1 mM Tris-HCl [pH 7.4], 130 mM NaCl, 5 mM KCl, 7.5 mM MgCl2) at ∼10 x the volume of the packed cells, and pelleted at 370 x g (at 4 °C). Pellets were washed a further two times in NKM buffer, prior to resuspension in 6 x volumes of homogenization buffer (10 mM Tris-HCl [pH 6.7], 10 mM KCl, 0.15 mM MgCl2, 1 mM DTT) Supplementarylemented with a protease inhibitor tablet, and incubated on ice for 10 mins. Cells were disrupted with 30 stokes of a Dounce homogenizer, 1 volume of 2 M sucrose was then added, and cellular debris and nuclear material was removed by centrifugation (1’200 x g, 5 mins, 4 °C). Lysates were clarified twice more by centrifugation. Mitochondria were pelleted at 7’000 x g for 10 mins and then resuspended in mitochondrial suspension buffer (10 mM Tris-HCl [pH 6.7], 1 mM DTT, 0.25 mM sucrose, 0.15 mM MgCl2) Supplementarylemented with a protease inhibitor tablet, and pelleted once more at 9’500 x g for 5 mins (4 °C). Extracts were produced by sonication of mitochondrial suspensions in (4 °C) SDS-PAGE sample buffer, and analysed by western blotting. Following western blotting, the relative change in abundance of PSKH2 in mitochondrial extracts was calculated by densitometry using ImageJ software. To account for variability in transfection efficiency between samples, all PSHK2 values were first normalised to the density of PSKH2 in the whole lysate, and then normalised to the mitochondrial marker ATP5A.

### Nuclear extraction

HEK-293T cells transfected with PSKH2-MYC or EGFP were washed in ice cold PBS 48 h post transfection, and resuspended in hypotonic lysis buffer (20 mM Tris-HCl [pH 7.4], 10 mM KCl, 2 mM MgCl2, 0.5 mM DTT, and 1 mM EDTA) supplemented with a protease inhibitor tablet, and incubated on ice for ∼5 mins. NP-40 was added to a final concentration of 1 % (v/v), the lysates were vortex mixed and incubated on ice for 3 mins. Cytoplasmic and nuclear fractions were separated by centrifugation at 800 x g for 8 mins (4 °C), and the cytoplasm containing supernatant was further clarified by centrifugation at 1’500 x g for 5 mins (4 °C). The pellet containing the nuclear fraction was washed briefly in hypotonic lysis buffer and proteins were extracted using SDS-sample buffer

### Metabolic labelling

HEK-293T cells were cultured and transfected as described above. 12 h post transfection, cells were washed in PBS and incubated with fresh DMEM medium supplemented with 100 µM alkynyl myristic acid for 18 h. Cells were lysed in DDM containing lysis buffer and PSKH2 was affinity precipitated as previously described. An on bead click chemistry reaction was accomplished using 100 µl Click-&-Go click chemistry reaction buffer kit and 40 µM azide-biotin as per the manufacturer’s instructions (Click Chemistry Tools), and incubated in the dark at room temperature for 30 mins with gentle agitation. PSKH2 containing affinity resin was then washed three times in 50 mM Tris-HCl (pH 7.4) and 150 mM NaCl and protein was eluted with SDS-PAGE sample buffer and analysed by western blotting.

### Molecular dynamics (MD) simulation of full-length PSKH2 and molecular modelling of the PSKH2/HSP90/Cdc37 complex

Unbiased full atom MD simulation of the full length AF2 model of human PSKH2 (Uniprot ID: Q96QS6) was performed for 700 nano-second (ns) using GROMACS 2020 [76]. Before using Amber99SB-ILDN [77] forcefield to parameterize the protein, hydrogen atoms were converted to virtual sites to remove the fastest vibrational freedom. The protein was further solvated with TIP3P waters in a dodecahedron box with charges neutralized by addition of sodium and chloride ions. The neighbour list for non-bonded interactions was defined using Verlet cutoff [78] [79] and Particle Mesh Ewald (PME) was used to calculate long-range interactions. Steepest-descent followed by conjugate descent (Fmax< 500kJmol^−1^nm^−1^.) energy minimization steps were used. The canonical ensemble was carried out by heating the system from 0 K to 310 K, using velocity rescaling for 100 ps [80]. The isothermal–isobaric ensemble (P = 1 bar, T = 310 K) was carried out using the Berendsen barostat for 100 ps. The unrestrained MD productions were collected using a time step of 5 fs. The trajectories were processed and analysed using the built-in GROMACS tools. Visualization was done using PyMOL 2.3.2. Locally installed version of the AlphaFold-Multimer [81] was used to model the complex between PSKH2, HSP90, and Cdc37.

## Supplementary Video 1

Movie of intrinsic fluctuations in full length PSKH2 derived from 700ns molecular dynamics simulations. See Supplementary Figure 9B for coloring scheme.

## Supplementary Video 2

Movie of PSKH2, HSP90A/B and Cdc37 complex generated from AlphaFold-Multimer. See Supplementary Figure 9C for coloring scheme.

## Data availability

All datasets generated for this current study are available from the corresponding authors on request and will be made available in https://fairsharing.org/IDGProject upon publication. MS data is available on ProteomeExchange.

## Competing Interests

The authors declare no competing interests.

## CRediT Contributions

Dominic P Byrne and Patrick A Eyers: Conceptualization, Formal analysis, Investigation, Methodology, Writing — original draft, Writing — review and editing. Leonard A Daly, Claire E Eyers, Safal Shrestha, Natarajan Kannan: Conceptualization, Formal analysis, Investigation, Methodology, Writing, review and editing. Vanessa Marensi: Investigation, Writing — review and editing; Krithika Ramakrishnan: Formal analysis, Writing — review and editing.

## Funding

N.K. and P.A.E. acknowledge funding from the NIH Illuminating the Druggable Genome (IDG) common fund consortium (NCI U01CA239106). D.P.B, K. R., L.A.D., P.A.E. and C.E.E. also acknowledge funding from BBSRC grants BB/S018514/1 and BB/N021703/1 and North West Cancer Research (NWCR) grants CR1097 and CR1208 (to V. M).

## Abbreviations

AF2: AlphaFold 2
DAVID: Database for Annotation,
Validation: and Integrated Discovery
EGFP: Enhanced Green Flourescent Protein
MS: mass spectrometry
PSKH1: Protein Serine Kinase
H1: PSKH2, Protein Serine Kinase H2.

## Supplementary Figures

**Supplementary Figure 1.**
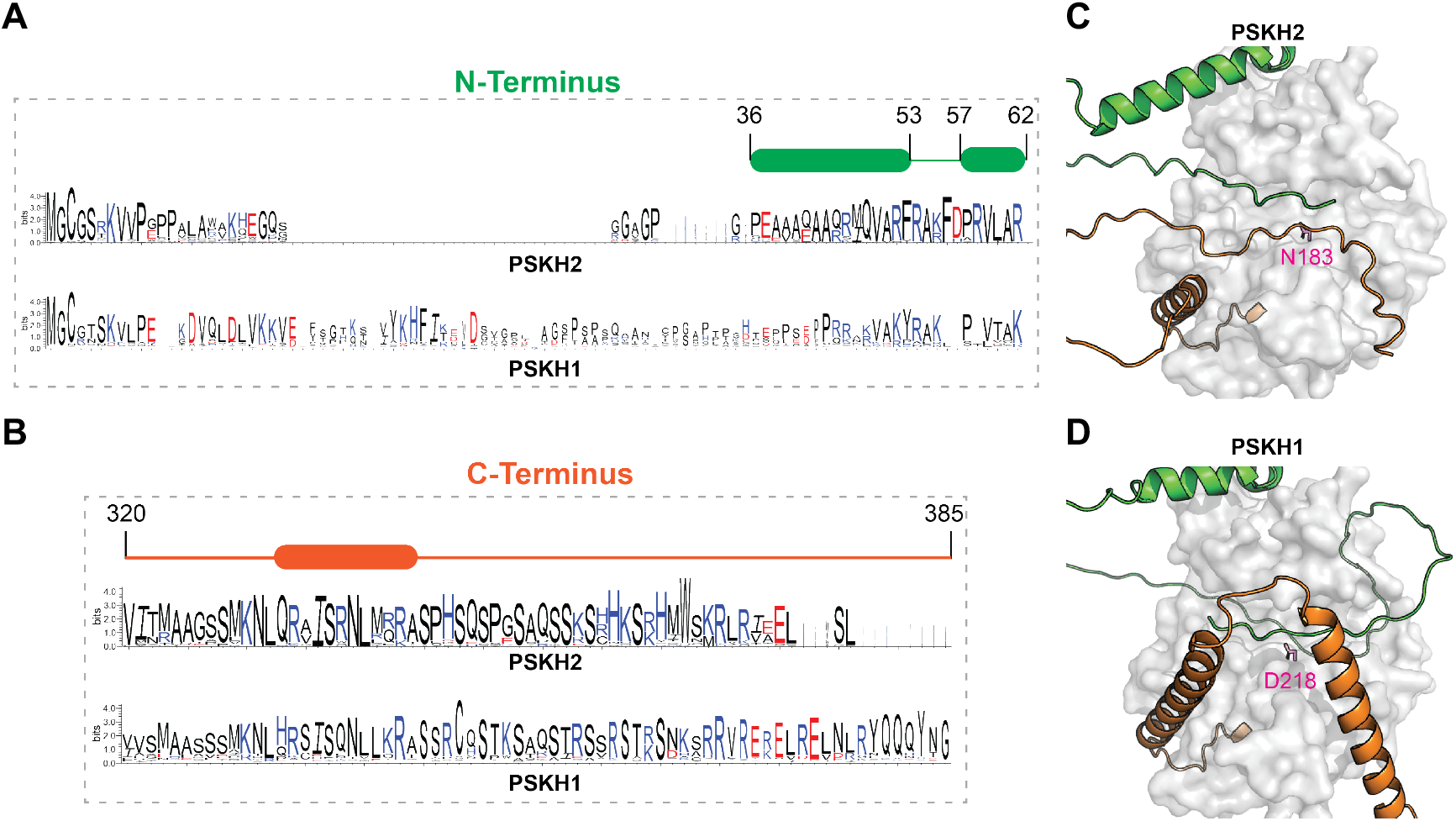
Sequence and structural basis of N- and C-terminal flanking region interactions with PSKH2 and PSKH1 kinase domains. **(A)** Weblogo [82] of the N-terminal flanking region within the PSKH2 and PSKH1 orthologs. The region corresponding to the predicted helices in the region is indicated. **(B)** Weblogo of the C-terminal flanking region of the PSKH2 and PSKH1 sequences. The region corresponding to the predicted helix is indicated. **(C)** AlphaFold model of human PSKH2 (Uniprot ID: Q96QS6) **(D)** AlphaFold model of human PSKH1 (Uniprot ID: P11801) **(C, D)** N and C-termini flanking regions are colored in green and orange respectively and represented as cartoon. Kinase domain is shown as surface and colored in gray. Catalytic residues HRD-Asp (D218) in PSKH1 and HRN-Asn (N183) are shown as sticks and colored in magenta.

**Supplementary Figure 2.**
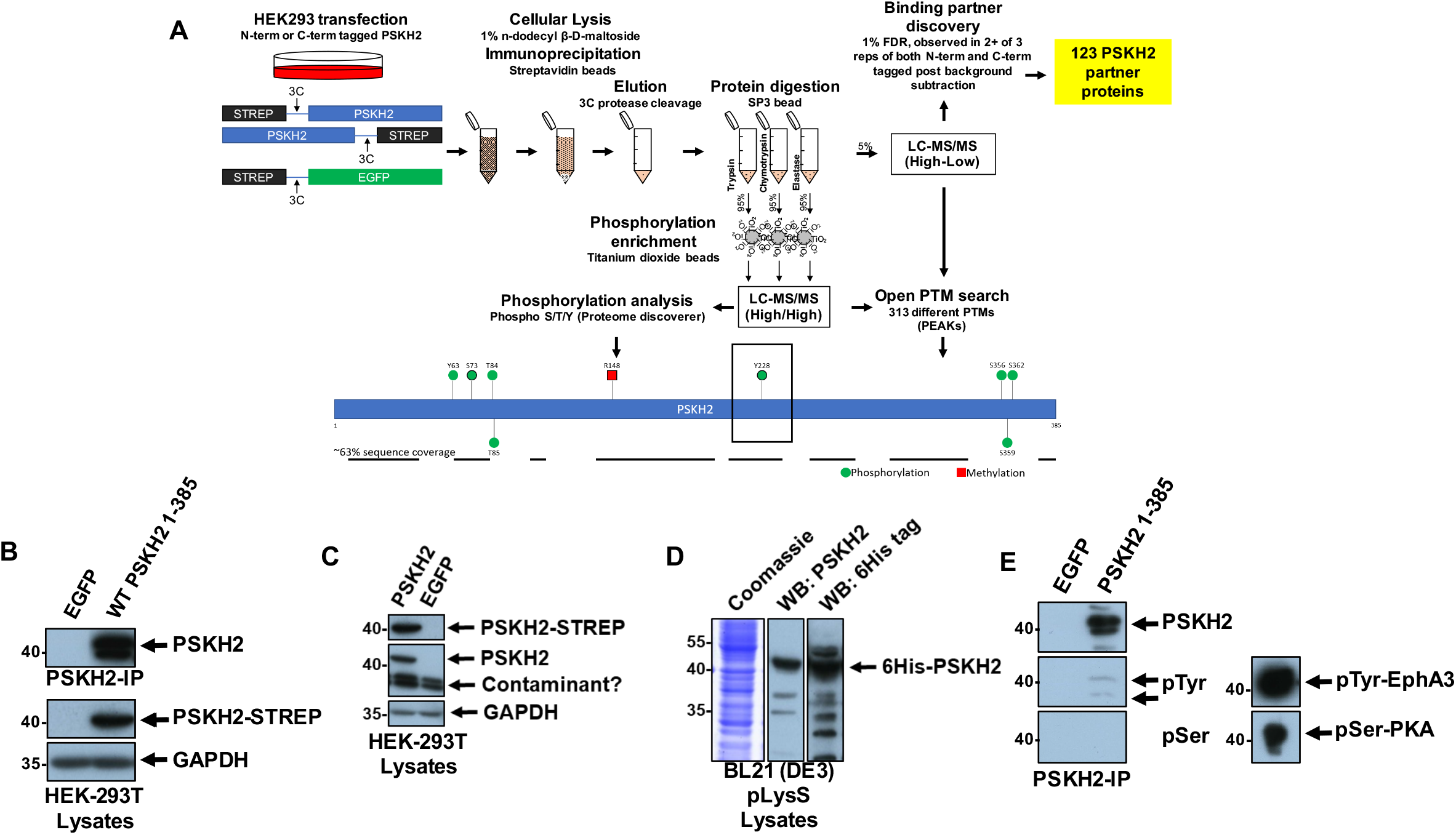
Immunoprecipitation of PSKH2 and evaluation of a PSKH2 polyclonal antibody. **(A)** Experimental transfection, IP and MS protocols and reported sites of PSKH2 modification (bottom). **(B)** Immunoprecipitation of exogenous CT-strep-tagged PSKH2 from HEK-293T cells. PSKH2 was detected with a non-commercial antibody with specificity towards PSKH2 in precipitates and an antibody directed towards tandem-strep tags in cell lysates. **(C, D)** Western blot showing overexpression of human PSKH2 in HEK-293T or IPTG-induced BL21 (DE3) pLysS *E*.*coli* cell extracts employing antibodies with specificity for Strep-tagged proteins, 6xHis-tagged proteins or PSKH2. **(E)** Immunoprecipitation of CT-Myc tagged PSKH2 from HEK-293T cells. Western blots show total PSKH2, and total phosphorylated Ser/Thr and Tyr. PKA and EphA3 are used as positive controls for Ser/Thr and Tyr phosphorylated proteins respectively, and were purified from or BL21 (DE3) pLysS *E*.*coli* as 6xHis tagged proteins.

**Supplementary Figure 3.**
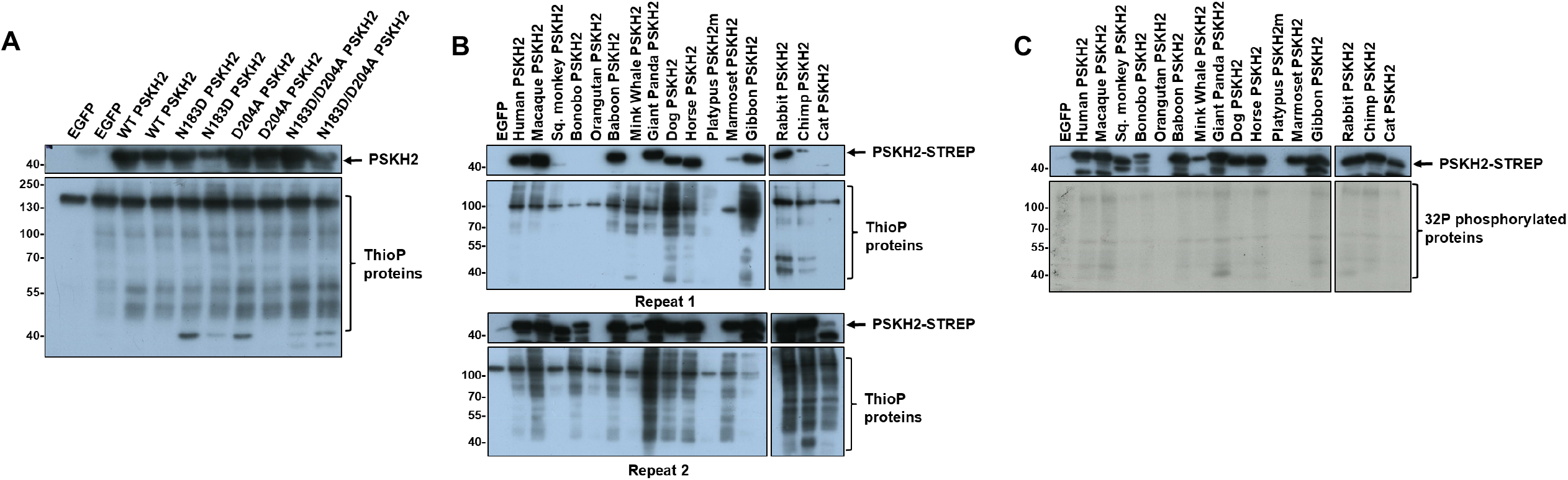
Evaluation of phosphotransferase activity. **(A)** Independent repeat of kinase assays using precipitated PSKH2 from HEK-293T cells. Protein phosphorylation as a consequence of kinase activity was detected using an antibody with specificity towards thiophosphate esters (Thio-P), and is compared with precipitations from cells overexpressing EGFP, WT PSKH2, or canonical kinase mutants of PSKH2. **(B)** Kinase activity in affinity precipitates of different species of PSKH2. Kinase assays were performed using ATP-γ-S **(B)** or **(C)** [γ-^32^P] ATP.

**Supplementary Figure 4.**
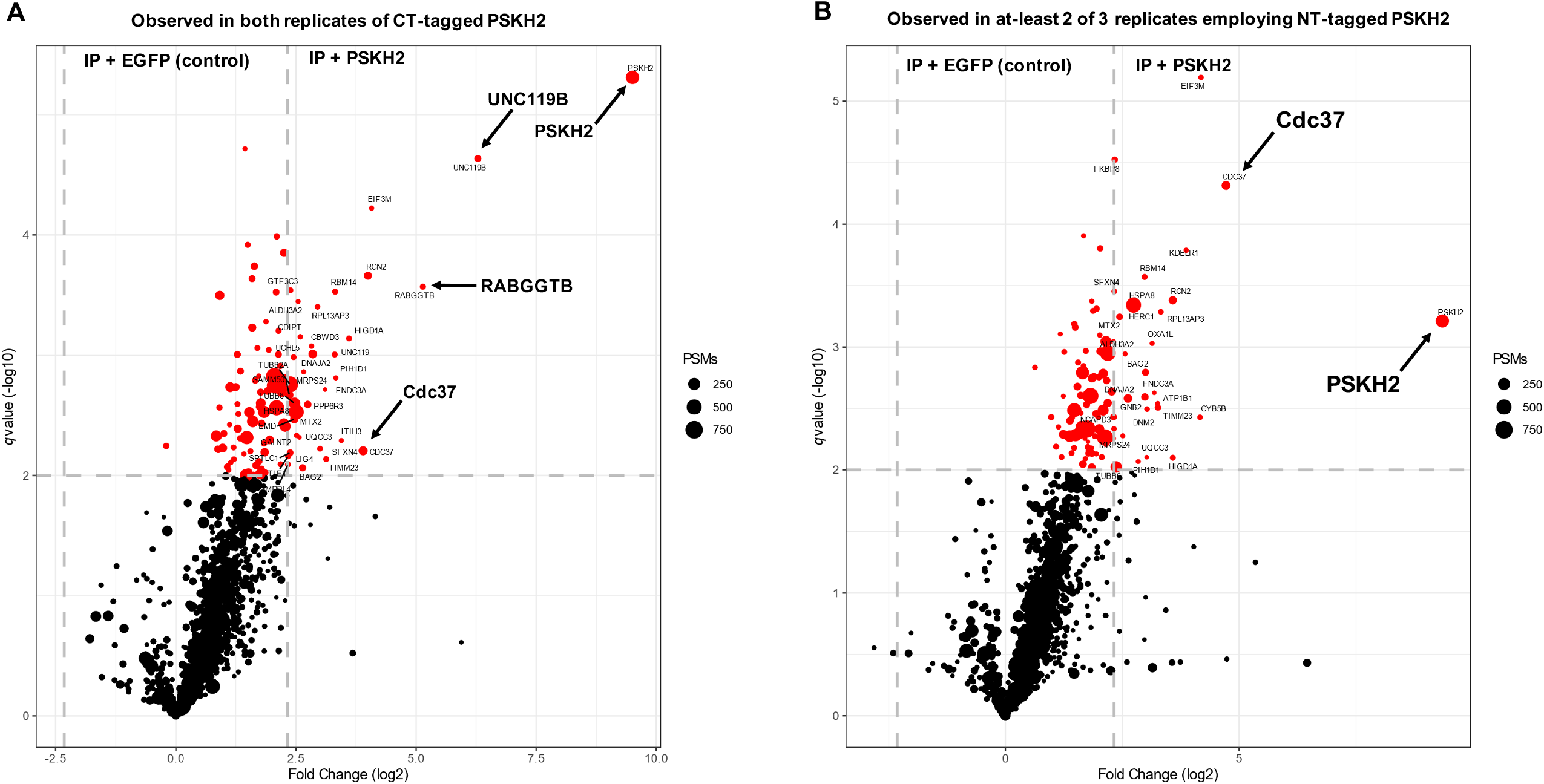
Analysis of the protein interactome of CT- and NT-tagged PSKH2. **(A)** Volcano plot depicting label-free protein quantification from PSKH2 IPs, shown as −log10(*q* value) versus log2 (1/21% abundance fold change). Dot size equates to the combined number of MS/MS events for a protein across all replicates. Black, *q* value > 0.01; red, *q* value < 0.01; significant proteins with a comparative fold change >5 are labelled with their gene name. Gray-dashed lines denote fivefold change and *q* value = 0.01. Data shown represents changes in abundance of proteins co-immunoprecipitated with CT-tagged PSKH2 and **(B)** NT-tagged PSKH2 compared to an EGFP-transfected control experiment. For both comparisons, only proteins observed in at least two replicate IPs were analysed.

**Supplementary Figure 5.**
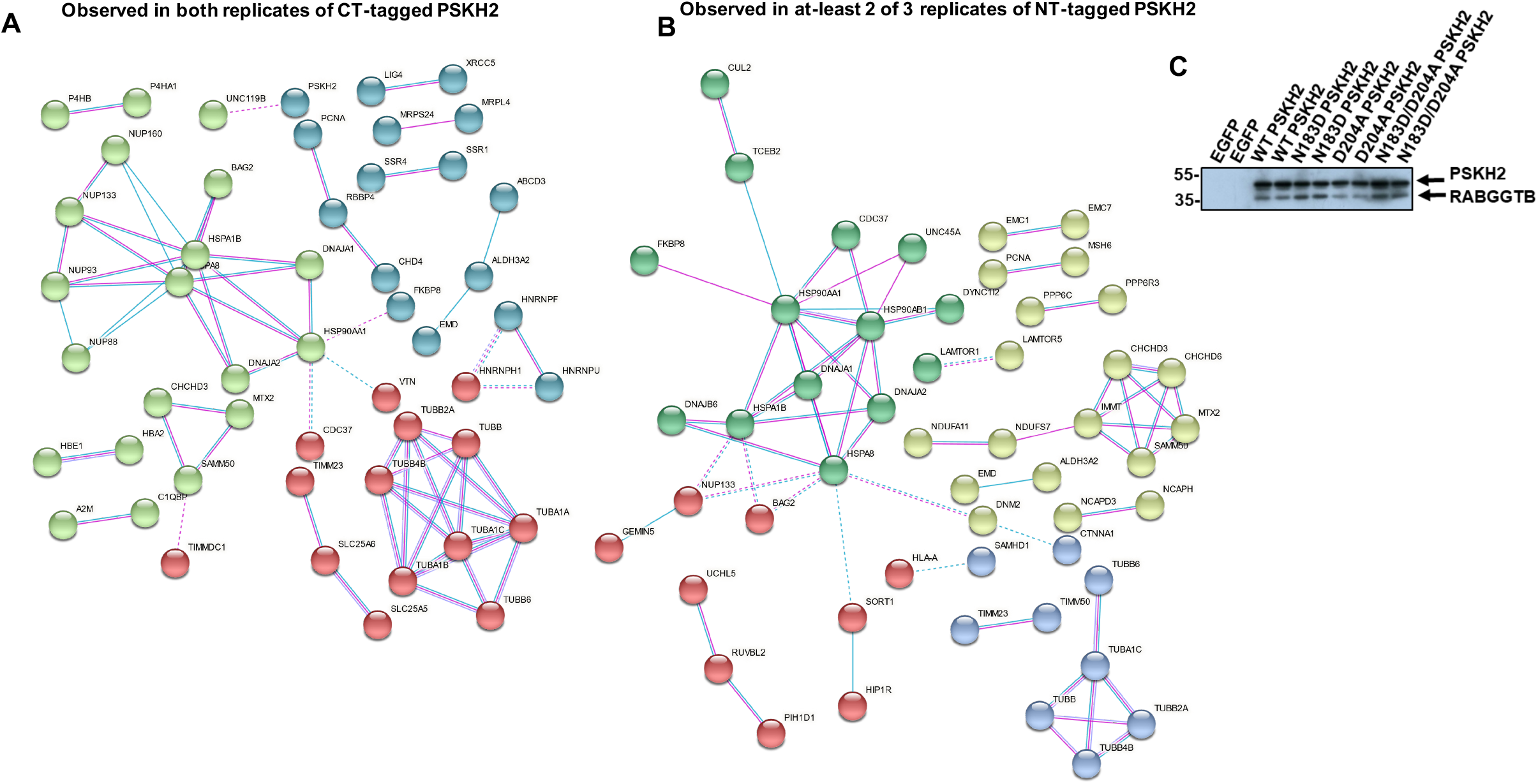
Reconstructed Protein-protein interaction networks present in CT- and NT-tagged PSKH2 immunoprecipitates. **(A)** Protein-protein interaction networks for significant binding partners (*p* value < 0.01) in both CT and **(B)** NT-tagged PSKH2 IPs.(C) Confirmation of RABGGTB binding.

**Supplementary Figure 6.**
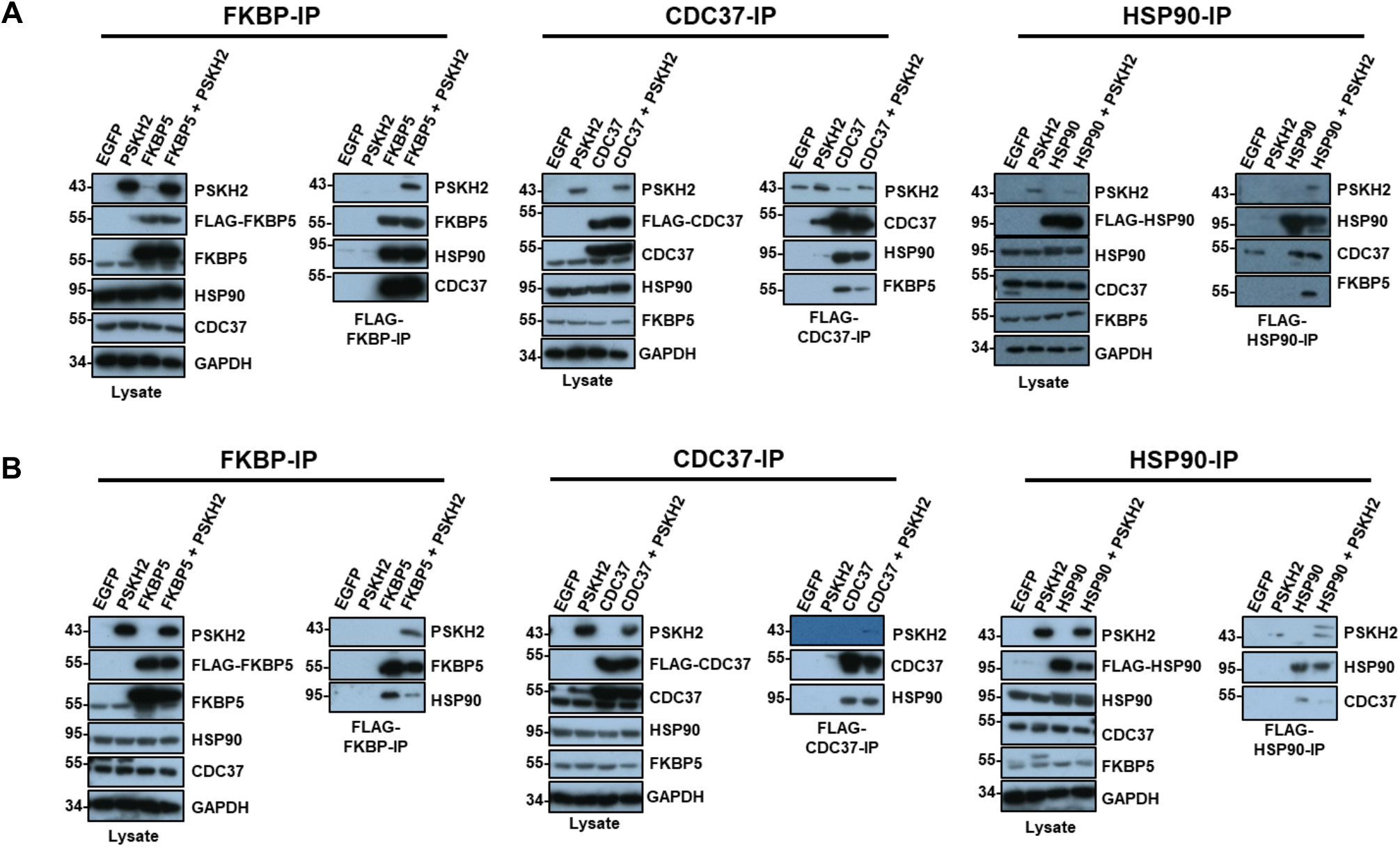
Co-immunoprecipitation of PSKH2 with FLAG-tagged HSP90 and co-chaperone proteins. PSKH2 (CT-Strep tag) and FLAG-tagged variants of the indicated HSP90 chaperone of co-chaperone proteins were expressed individually or as a co-expression in HEK-293T cells. FLAG-tagged HSP90 proteins or co-chaperones were immunoprecipitated and blotted for the indicated proteins. Results from two independent experiments are shown (top and bottom).

**Supplementary Figure 7.**
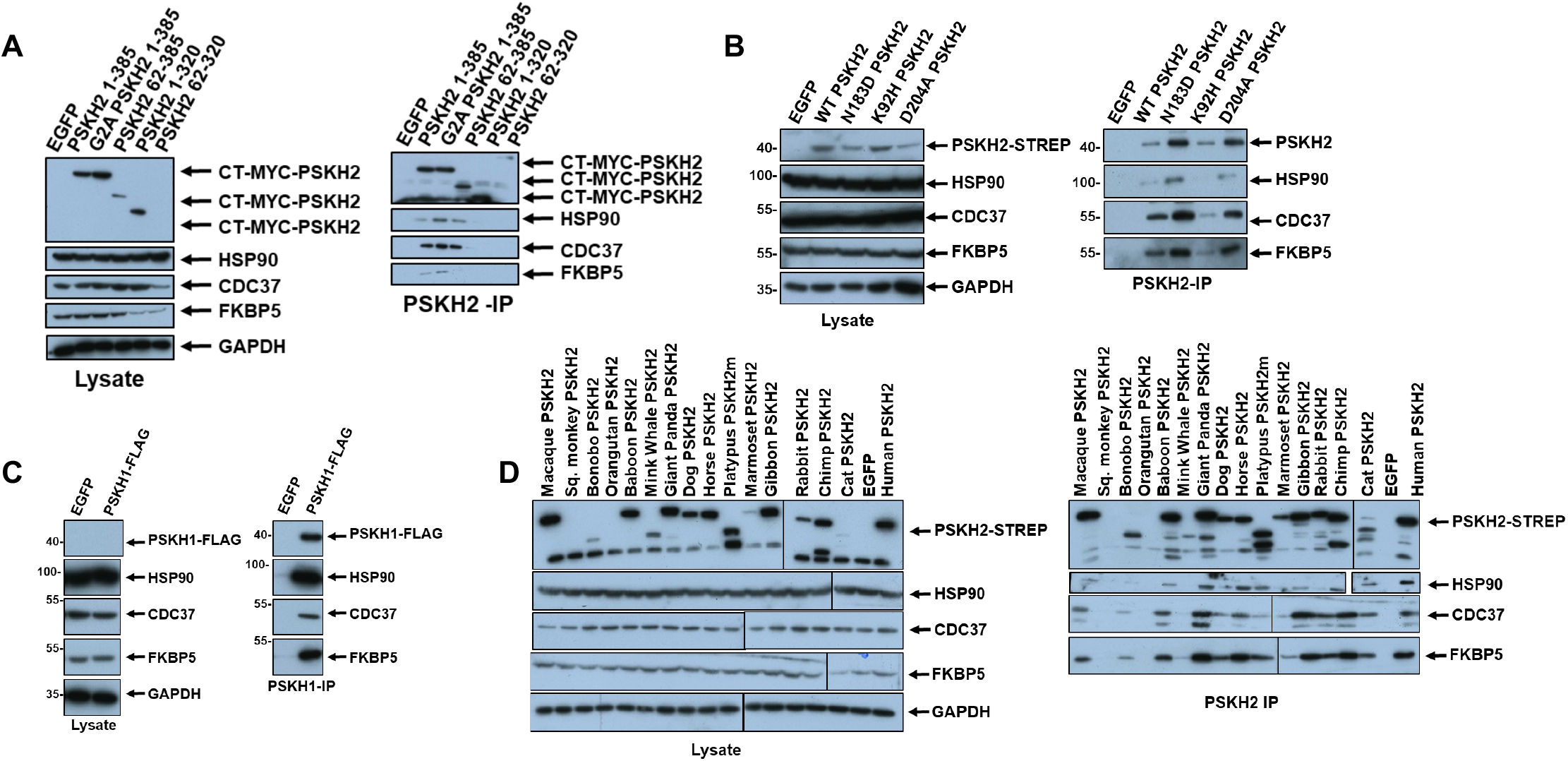
PSKH2 interacts with HSP90 and co-chaperones. **(A)** Independent repeat showing co-precipitation of HSP90 proteins with full-length and truncated PSKH2 constructs. **(B)** Independent repeat showing co-precipitation of HSP90 proteins with WT and canonical kinase activity mutants of PSKH2. **(C)** Co-precipitation of HSP90 proteins with PSKH1. **(D)** Co-precipitation of HSP90 proteins with different species of PSKH2.

**Supplementary Figure 8.**
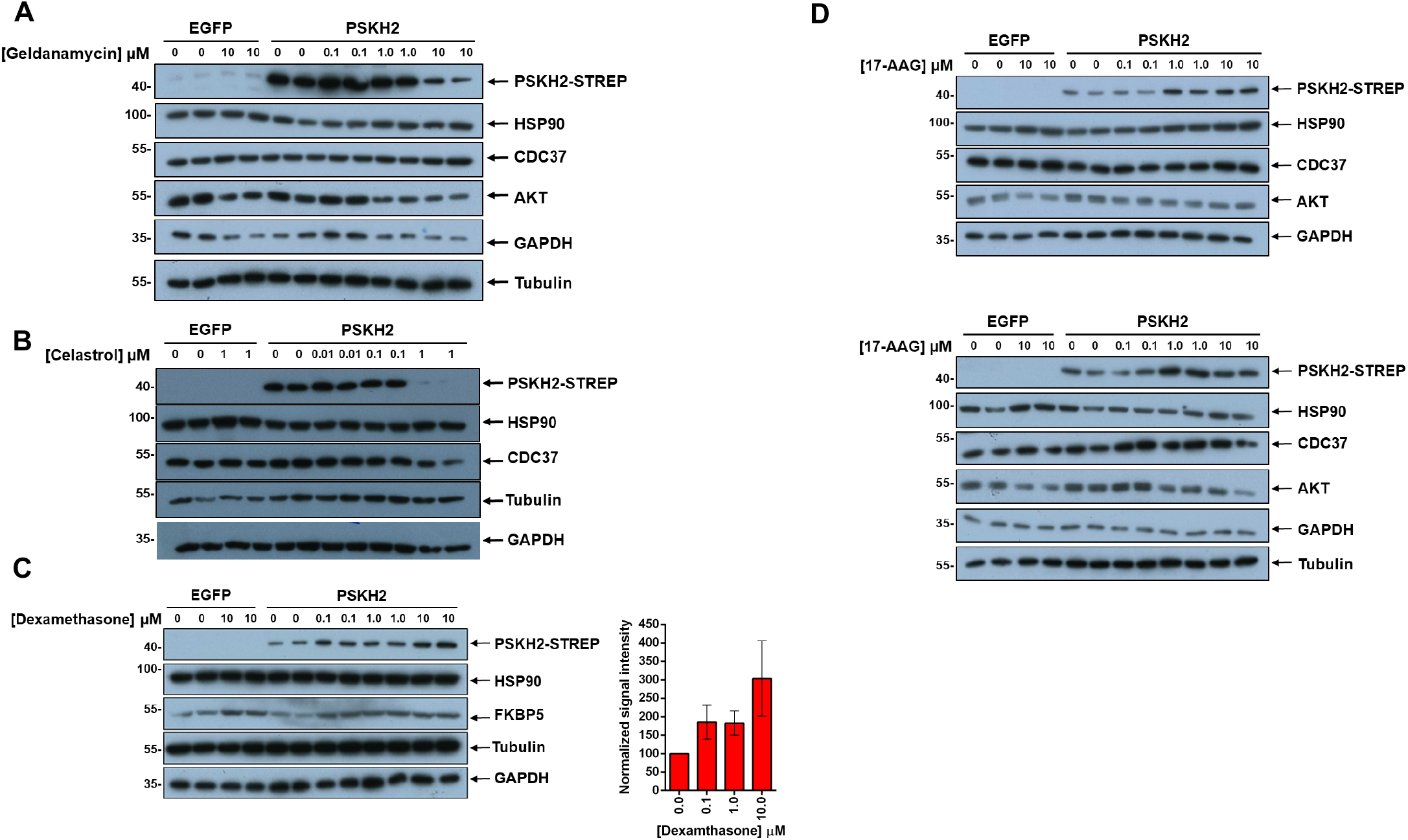
PSKH2 expression is controlled by HSP90. **(A)** Independent repeat showing reduction of PSKH2 expression in HEK-293T cells with increasing concentration of geldanamycin and **(B)** celasterol. **(C)** Western blot showing increased expression of PSKH2 and FKBP5 in the presence of dexamethasone. Right panel, densitometry data shown is mean and SD of PSKH2 band intensity from four independent experiments, first normalized to tubulin and then PSKH2 bands corresponding to control (DMSO) conditions. **(D)** Two independent repeats (top and bottom panel) comparing PSKH2 expression in the presence of 17-AAG.

**Supplementary Figure 9.**
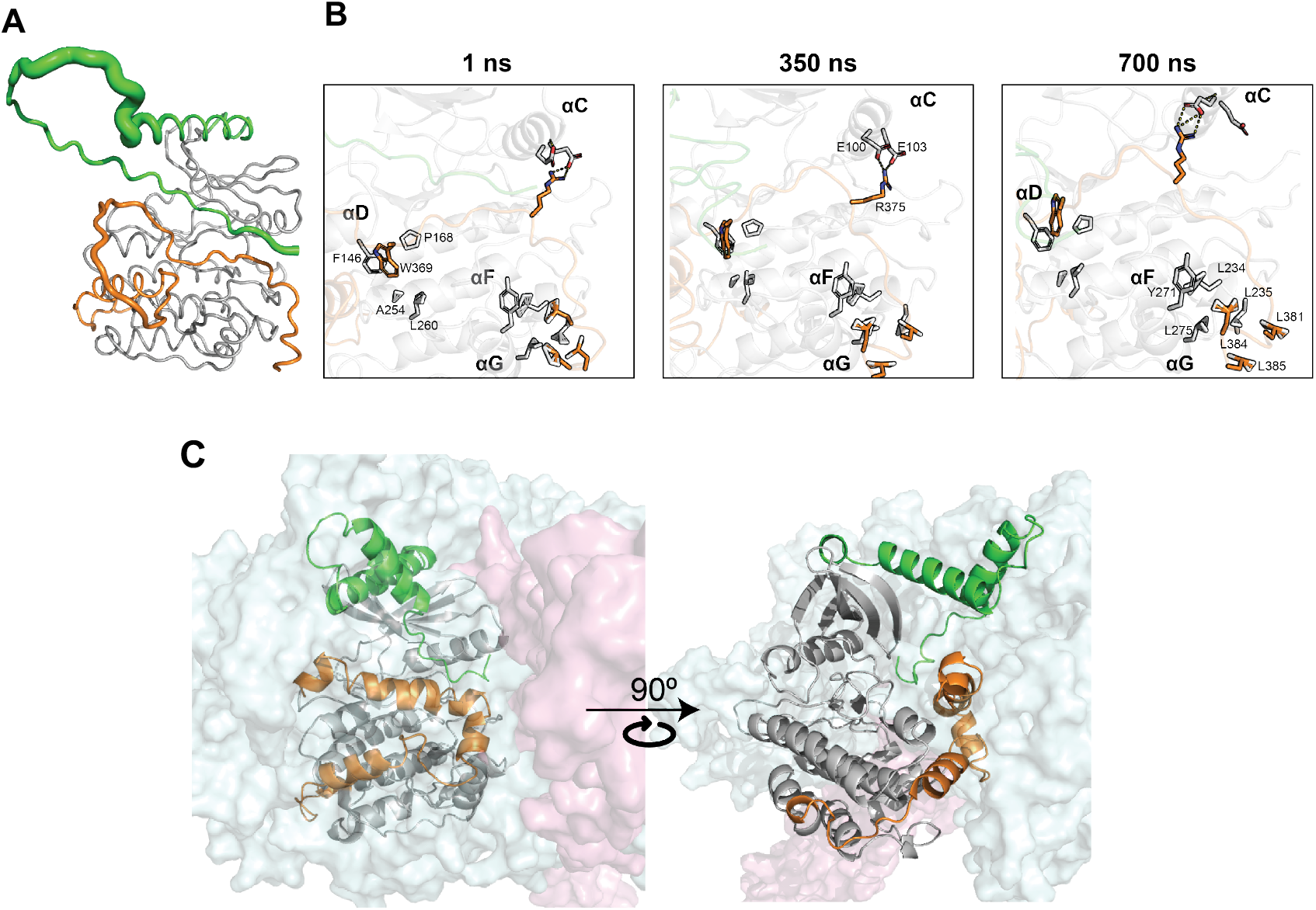
Molecular dynamics (MD) simulation of Full length human PSKH2 and modelling of its binding mode to the HSP90/Cdc37 complex. (**A**) Putty representation of the full length human PSKH2 summarizing the Root Mean Square Fluctuations (RMSF) during the 700 ns Molecular Dynamics (MD) simulation (**B**) Snapshots of the MD simulation of full length PSKH2 showing hydrophobic interactions and salt bridges formed by the C-terminal flanking region with the pseudokinase domain. (**C**) AlphaFold-multimer model of full length human PSKH2, HSP90 A/B (Uniprot IDs: P07900/P08238) and CDC37 (Uniprot ID: Q16543) complex. HSP90 and Cdc37 are shown in surface representation and colored in cyan and magenta respectively. (**B**,**C**) PSKH2 is shown as cartoon with pseudokinase domain, N-terminal, and C-terminal regions colored in gray, green, and orange, respectively.

**Supplementary Figure 10.**
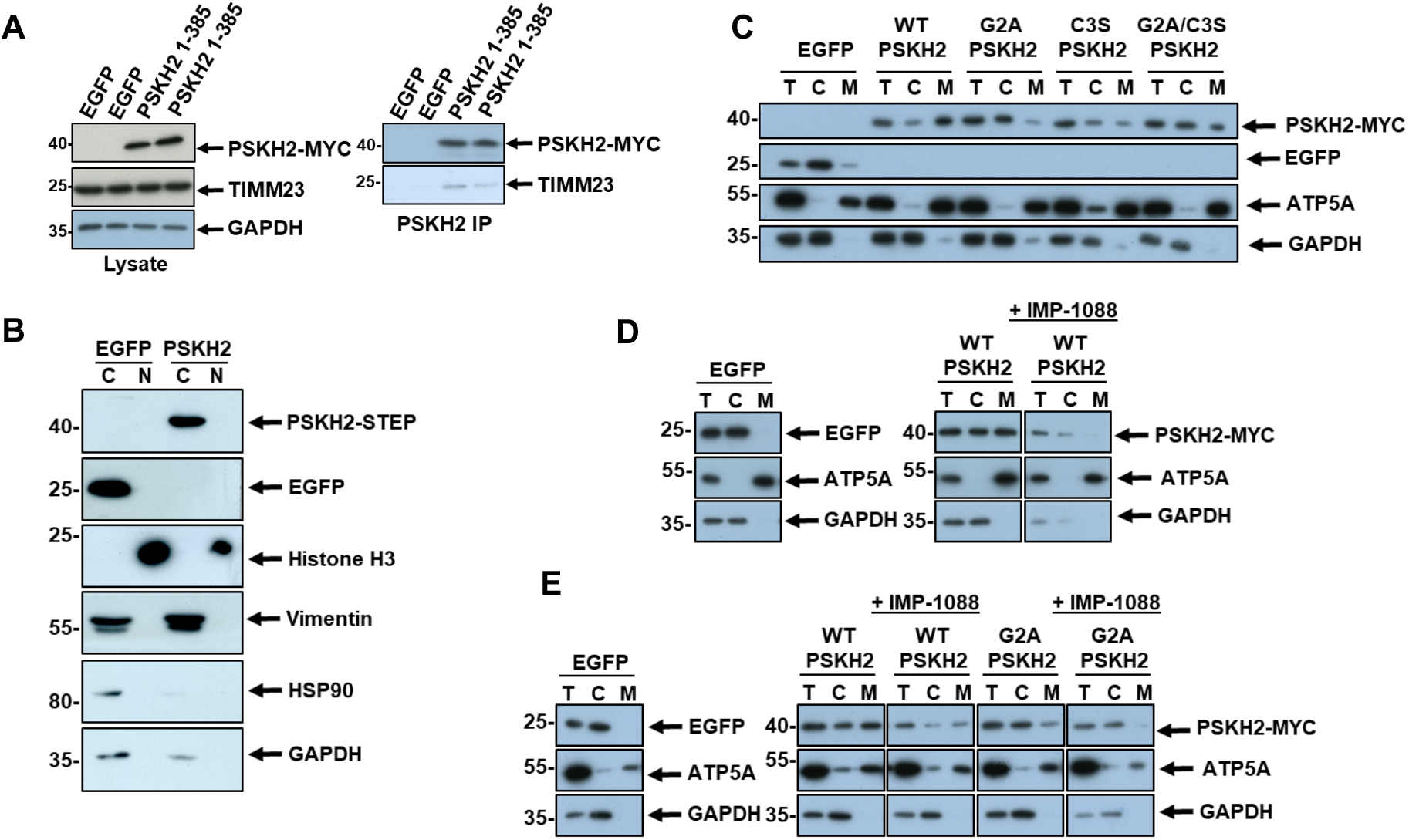
Incorporation of PSKH2 in mitochondrial-enriched fractions is dependent on N-myristoylation. **(A)** Independent repeat showing co-precipitation of mitochondrial protein TIMM23 with PSKH2. **(B)** Nuclear fractionation of HEK-293T cells overexpressing EGFP of CT-Myc PSKH2. **(C)** Independent repeat showing mitochondrial fractionation of HEK-293T cells overexpressing EGFP, WT PSKH2 (CT-Myc) or PSKH2 mutated at putative sites of acylation. **(D-E)** Independent repeats showing mitochondrial fractionation of HEK-293T treated in the presence or absence of IMP-1088. For all mitochondrial fractionations; T = total cell lysate, C = cytoplasmic fraction, M = mitochondrial fraction.

## Notes

### Competing Interest Statement

The authors have declared no competing interest.

### Summary of Updates

Inclusion of additional references omitted from previous version.

